# Renal medullary carcinomas depend upon *SMARCB1* loss and are sensitive to proteasome inhibition

**DOI:** 10.1101/487579

**Authors:** Andrew L. Hong, Yuen-Yi Tseng, Jeremiah Wala, Won Jun Kim, Bryan D. Kynnap, Mihir B. Doshi, Guillaume Kugener, Gabriel J. Sandoval, Thomas P. Howard, Ji Li, Xiaoping Yang, Michelle Tillgren, Mahmoud Ghandi, Abeer Sayeed, Rebecca Deasy, Abigail Ward, Brian McSteen, Katherine M. Labella, Paula Keskula, Adam Tracy, Cora Connor, Catherine M. Clinton, Alanna J. Church, Brian D. Crompton, Katherine A. Janeway, Barbara Van Hare, David Sandak, Ole Gjoerup, Pratiti Bandopadhayay, Paul A. Clemons, Stuart L. Schreiber, David E. Root, Prafulla C. Gokhale, Susan N. Chi, Elizabeth A. Mullen, Charles W. M. Roberts, Cigall Kadoch, Rameen Beroukhim, Keith L. Ligon, Jesse S. Boehm, William C. Hahn

**Affiliations:** Boston Children’s Hospital, 300 Longwood Avenue, Boston, Massachusetts, 02115, USA; Dana-Farber Cancer Institute, 450 Brookline Avenue, Boston, Massachusetts, 02215 USA; Broad Institute of Harvard and MIT, 415 Main Street, Cambridge, Massachusetts, 02142 USA; Rare Cancer Research Foundation, 112 S. Duke Street, Suite 101, Durham, NC 27701 USA; RMC Support, P.O. Box 73156, North Charleston, S.C. 29415 USA; St. Jude Children’s Research Hospital, St. Jude Children’s Research Hospital, 262 Danny Thomas Place, Memphis, Tennessee 38120, USA; Brigham and Women’s Hospital, 75 Francis Street, Boston, Massachusetts, 02115 USA

**Author notes:** **To whom correspondence should be addressed**: William C. Hahn, M.D., Ph.D., Department of Medical Oncology, Dana-Farber Cancer Institute, 450 Brookline Avenue, Dana 1538, Boston, MA 02215.

## Abstract

Renal medullary carcinoma (RMC) is a rare and deadly kidney cancer in patients of African descent with sickle cell trait. Through direct-to-patient outreach, we developed genomically faithful patient-derived models of RMC. Using whole genome sequencing, we identified intronic fusion events in one *SMARCB1* allele with concurrent loss of the other allele, confirming that SMARCB1 loss occurs in RMC. Biochemical and functional characterization of these RMC models revealed that RMC depends on the loss of *SMARCB1* for survival and functionally resemble other cancers that harbor loss of *SMARCB1*, such as malignant rhabdoid tumors or atypical teratoid rhabdoid tumors. We performed RNAi and CRISPR-Cas9 loss of function genetic screens and a small-molecule screen and identified *UBE2C* as an essential gene in SMARCB1 deficient cancers. We found that the ubiquitin-proteasome pathway was essential for the survival of SMARCB1 deficient cancers *in vitro* and *in vivo.* Genetic or pharmacologic inhibition of this pathway leads to G2/M arrest due to constitutive accumulation of cyclin B1. These observations identify a synthetic lethal relationship that may serve as a therapeutic approach for patients with SMARCB1 deficient cancers.

## Introduction

Renal medullary carcinoma (RMC) was first identified in 1995 and is described as the seventh nephropathy of sickle cell disease (Jr et al., 1995). RMC is a rare cancer that occurs primarily in patients of African descent that carry sickle cell trait and presents during adolescence with symptoms of abdominal pain, hematuria, weight loss and widely metastatic disease. Due to the aggressive behavior of this disease and the small numbers of patients, no standard of care exists, and patients are generally treated with therapies including nephrectomy, chemotherapy and radiation therapy. Despite, this aggressive regimen, the mean overall survival rate is only 6-8 months (Alvarez et al., 2015; Beckermann et al., 2017; Ezekian et al., 2017; Iacovelli et al., 2015). Gene expression of tumor samples from patients with RMC indicates that this entity is distinct from renal cell carcinomas and urothelial carcinomas (Swartz et al., 2002; Yang et al., 2004).

Recent studies have identified that SMARCB1 is lost at the protein level in RMC (Cheng et al., 2008). SMARCB1 is a tumor suppressor and core member of the SWI/SNF complex implicated in malignant rhabdoid tumors (MRT) and atypical teratoid rhabdoid tumors (ATRT)(Roberts et al., 2002). MRTs and ATRTs harbor few somatic genetic alterations and occur in young children (Gröbner et al., 2018; Lawrence et al., 2013; Lee et al., 2012; Ma et al., 2018). In contrast to MRTs and ATRTs, where there is biallelic loss of *SMARCB1* in the genome (Chun et al., 2016; Torchia et al., 2016), in RMCs, fusion events in *SMARCB1* have been identified in 4 of 5 patients (Calderaro et al., 2016) and 8 of 10 patients (Carlo et al., 2017) using whole exome sequencing (WES), fluorescence *in situ* hybridization (FISH), array comparative genomic hybridization (CGH), and RNA-sequencing. An unresolved question is whether these cancers depend upon loss of this tumor suppressor, since deletion of *SMARCB1* in mice does not lead to renal cancers (Roberts et al., 2002).

To date, limited tissue from these patients has prevented full characterization of RMC. Here, we characterized new patient-derived cell lines developed from samples following neoadjuvant therapy and at relapse. These cell lines have allowed us to functionally demonstrate dependence of RMC on loss of SMARCB1 for survival and to uncover the proteasome as a core druggable vulnerability.

## Results

### Derivation and genomic characterization of RMC models

We developed models from two patients with a diagnosis of RMC (**Methods**). For the first patient, we obtained the primary tissue from our local institution at the time of the initial nephrectomy. We generated a short-term culture normal cell line, CLF_PEDS0005_N, and a tumor cell line, CLF_PEDS0005_T1 (**Supp Fig. 1A**). In addition, we obtained fluid from a thoracentesis performed when the patient relapsed 8 months into therapy. We isolated two cell lines that grew either as an adherent monolayer, CLF_PEDS0005_T2A, or in suspension, CLF_PEDS0005_T2B. Each of these tumor cell lines expressed the epithelial marker, CAM5.2, and lacked expression of SMARCB1 similar to that observed in the primary tumor (**Supp Fig. 1B**). For the second patient, we partnered with the Rare Cancer Research Foundation and obtained samples through a direct-to-patient portal (www.pattern.org). The primary tumor tissue from the second patient was obtained at the time of the initial nephrectomy. From this sample, we generated the tumor cell line, CLF_PEDS9001_T1. Both patients received 4-8 weeks neoadjuvant chemotherapy prior to their nephrectomy.

Prior studies have identified deletion of one allele of *SMARCB1* along with fusion events in *SMARCB1* in RMC patients (Calderaro et al., 2016; Carlo et al., 2017). Specifically, one study identified *SMARCB1* fusion events in 8 of 10 patients utilizing FISH, while another study identified fusion events in 4 of 5 patients utilizing a combination of FISH, RNA-sequencing and Sanger sequencing. We performed WES (CLF_PEDS0005) or PCR-free whole genome sequencing (WGS; CLF_PEDS9001) on the primary kidney tumor tissue. In both patients, we found tumor purity was <20% and confirmed the presence of sickle cell trait (**Supp Fig. 1C**). This low tumor purity is attributable to the stromal desmoplasia seen in RMC (Swartz et al., 2002). Furthermore, we failed to identify homozygous or heterozygous deletions or mutations of *SMARCB1* due to the low tumor purity.

We performed WES on the normal cell line (CLF_PEDS0005_N) or whole blood (CLF_PEDS9001) and compared it to the primary tumor cell lines (CLF_PEDS0005_T1 and CLF_PEDS9001_T) and metastatic cell lines (CLF_PEDS0005_T2A and CLF_PEDS0005_T2B). We used the Genome Analysis Toolkit (GATK) v4.0.4.0 for variant discovery and copy number analyses (**Methods**) (McKenna et al., 2010) and found a low mutation frequency (1-3 mutations/mb; **Fig. 1A**) in the tumor cell lines similar to that of other pediatric cancers and cell lines such as MRT, ATRT and Ewing sarcoma (Cibulskis et al., 2013; Johann et al., 2016; Wala et al., 2018). We performed MuTect 2.0, a method to identify somatic point mutations, on these samples and filtered mutations based on the Catalogue of Somatic Mutations in Cancer (COSMIC) and found that only the metastatic cell lines harbored mutations in *TP53* and *TPR* (**Supp Data 1**) (Cibulskis et al., 2013). From our copy number analysis, we confirmed the heterozygous loss of *SMARCB1.* In agreement with prior studies, we failed to find an identifiable mutation or deletion to account for the loss of the second *SMARCB1* allele with WES.

**Fig. 1:**
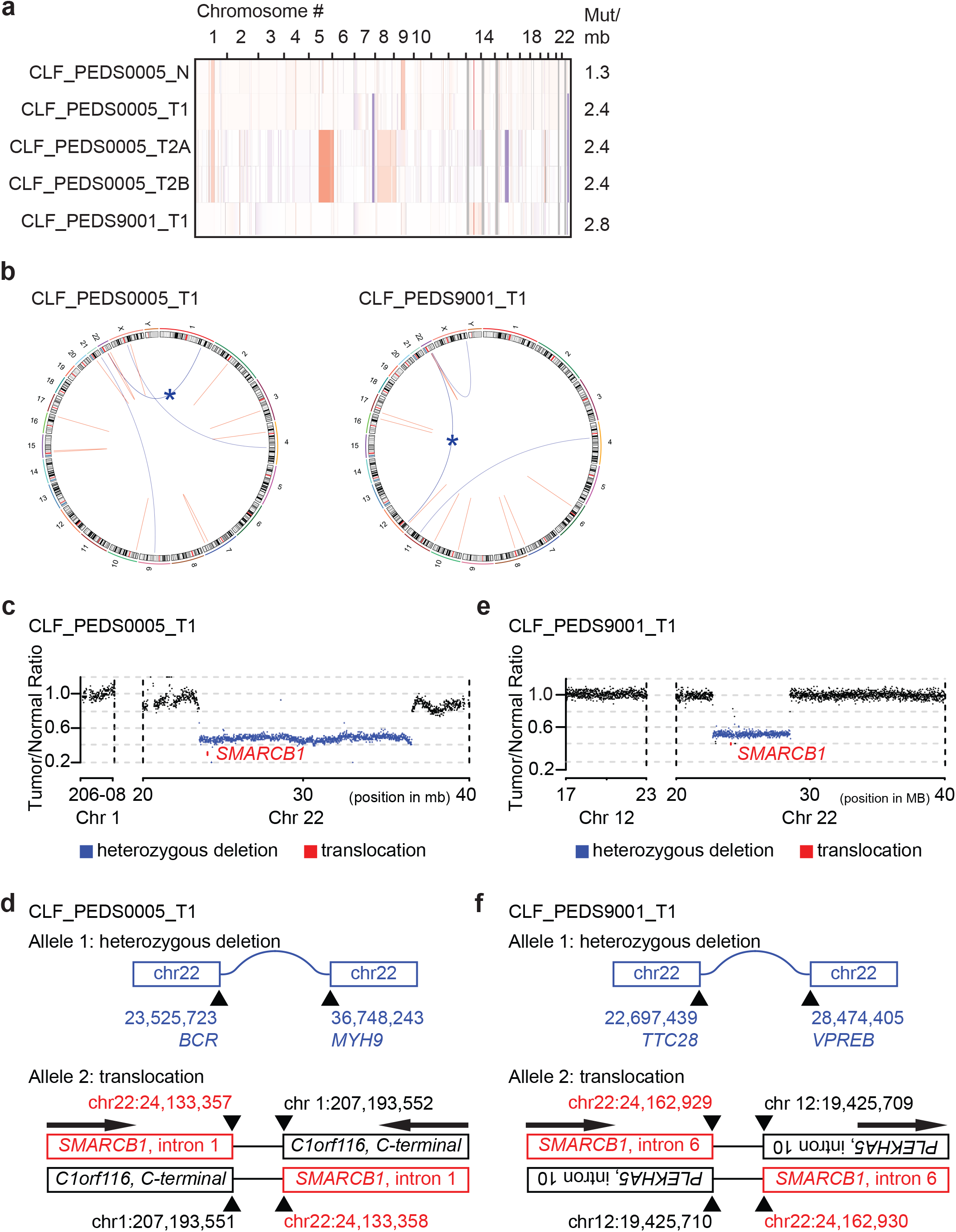
Patient derived models of RMC have quiet genomes and are driven by intronic translocations. **(a)** Copy number analysis of RMC models identifies heterozygous loss of alleles or low-level gains in primary and metastatic tumors. Rates of mutations per megabase are consistent with patients with RMC or other pediatric cancers such as rhabdoid tumor. **(b)** Circos plots of RMC cell lines. Red indicates a deletion. All deletions are located in the introns. Blue arcs indicate fusions identified with SvABA v0.2.1(Wala et al., 2018). Blue star indicates a *SMARCB1* rearrangement. **(c)** Read counts from PCR-free Whole Genome Sequencing identify single copy deletion of *SMARCB1* in CLF_PEDS0005_T1. **(d)** Second allele of *SMARCB1* is lost by a balanced translocation occurring in intron 1 of *SMARCB1* and fuses to the C-terminal end of C1orf116 in chromosome 1 in CLF_PEDS0005_T1. **(e)** Read counts from PCR-free Whole Genome Sequencing identify single copy deletion of *SMARCB1* in CLF_PEDS9001_T1. **(f)** Second allele of *SMARCB1* is lost by a balanced translocation occurring in intron 6 of *SMARCB1* and fuses to the anti-sense intron 10 of PLEKHA5 in chromosome 12 in CLF_PEDS9001_T1.

We then developed a dual-color break apart FISH using bacterial artificial chromosome (BAC) probes surrounding *SMARCB1* to ascertain if the loss of the second *SMARCB1* allele was from a fusion event as seen in prior studies (**Methods**) (Calderaro et al., 2016; Carlo et al., 2017). 83% of NA10851 B-lymphocyte control cells harbored 2 fused signals indicating the presence of two *SMARCB1* alleles. In contrast, 82% of cells from CLF_PEDS0005_T1 and 84% of CLF_PEDS9001_T exhibited only one signal and a faint fused signal. These observations suggest that the *SMARCB1* loci in these patient derived cell lines harbor one rearranged copy of *SMARCB1* resulting in one signal and a deletion of *SMARCB1* in the other allele resulting in the loss of the majority of the fused signal (**Supp Data 2**).

We subsequently performed PCR-free WGS to assess for structural variations that would not be captured by WES to elucidate the breakpoint of the rearrangement in *SMARCB1.* We achieved an average depth of coverage of 38x for the germline DNA and 69x for the tumor cell line DNA. We performed Structural variation and indel Analysis By Assembly (SvABA) v0.2.1 on these samples (Wala et al., 2018). In both patient-derived cell lines, we found several small indels in introns and translocations (**Fig. 1B; Supp Data 3-4**). For CLF_PEDS0005_T1, we found a large deletion between *BCR* and *MYH9* which is predicted to lead to loss of one allele of *SMARCB1* (**Fig. 1C and D**). We then identified a balanced translocation affecting *SMARCB1* that is predicted to disrupt SMARCB1 function. Specifically, we identified a fusion in intron 1 of *SMARCB1* to the intron region following the C-terminal end of *C1orf116*, yielding a non-functional allele. For CLF_PEDS9001_T1, we identified a large deletion between *TTC28* and *VPREB* that would lead to loss of one allele of *SMARCB1* (**Fig. 1E and F**). We then identified a balanced translocation that leads to fusion of intron 10 of *PLEKHA5* to intron 6 of *SMARCB1.* These findings confirmed prior work that identified deletion of one allele and fusions involving the other allele of *SMARCB1* in RMC (Calderaro et al., 2016; Carlo et al., 2017).

We then performed qRT-PCR to confirm that *SMARCB1* was lost at the introns identified by WGS in our cell lines. Specifically, we interrogated each exon-exon junction of *SMARCB1* by developing primers that spanned each junction. We generated cDNA libraries from TC32, an Ewing Sarcoma *SMARCB1* wild-type cell line, and from our RMC models. As compared with TC32, we failed to find exon-exon cDNA in CLF_PEDS0005_T1 suggesting that the translocation in *SMARCB1* occurred between exons 1-2 (**Supp Fig. 1D**). For CLF_PEDS9001_T, we found that *SMARCB1* expression was approximately 50% of that observed in the wild type *SMARCB1*-expressing cell line TC32 for the first 5 exons, but that none of the exons were expressed after exon 5. This observation confirmed that the translocation in *SMARCB1* in CLF_PEDS9001_T occurs in intron 6 of *SMARCB1* (**Supp Fig. 1D; Methods**). We then assessed the genomic DNA of the primary tumor tissues and designed primers surrounding the fusions identified by WGS. We confirmed that the fusions identified in the cell lines could be identified in the genomic DNA of the primary tumor tissues (**Supp Fig. 1E-G; Methods**). Taken together, we have developed models from two patients with RMC which faithfully recapitulate known genomics of this disease.

### Patient-derived models of RMC are similar to SMARCB1 deficient cancers

We performed RNA-sequencing and transcriptomic profiling to compare the RMC models to other renal tumors or tumors that harbor loss of *SMARCB1.* Specifically, we compared the Therapeutically Applicable Research to Generate Effective Treatments (TARGET) RNA-sequencing data from pediatric renal tumors (e.g. Wilms Tumor, Clear Cell Sarcoma of the Kidney, and Malignant Rhabdoid Tumor) or normal kidney tissues with the RMC models using t-distributed stochastic neighbor embedding (tSNE) (**Methods**). The normal cell line, CLF_PEDS0005_N, clustered with TARGET normal kidney tissues and RMC cell lines from both patients clustered with the TARGET Rhabdoid Tumor samples (**Fig. 2A**). These observations showed that these RMC cell lines share expression patterns with patients with MRTs.

**Fig. 2:**
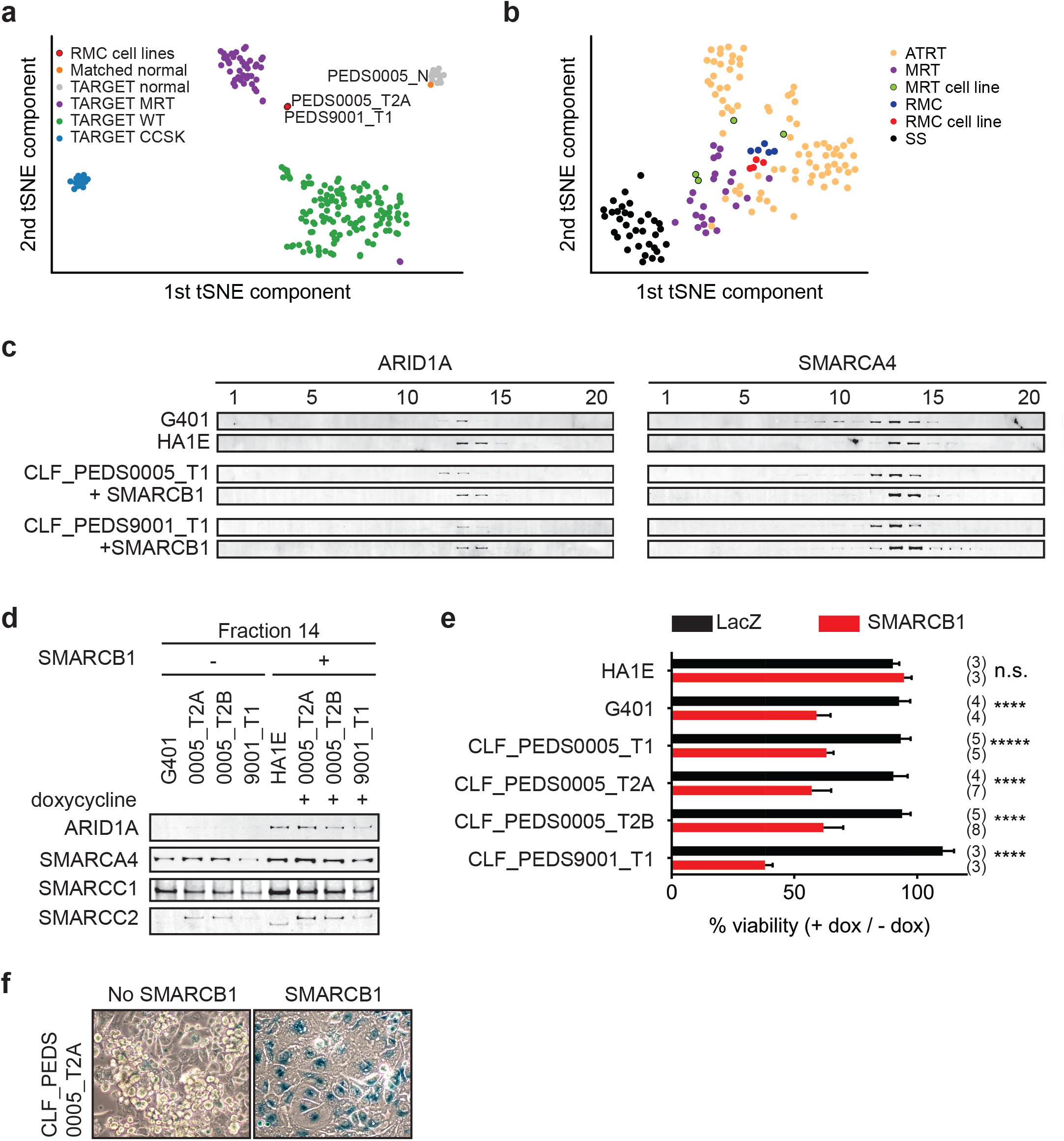
Patient derived models of RMC are functionally similar to SMARCB1 deficient cancers. **(a)** Utilizing tSNE, gene expression by RNA-sequencing from patients with pediatric renal tumors profiled in TARGET such as clear cell sarcoma of the kidney (CCSK), malignant rhabdoid tumor (MRT) or Wilms tumor (WT) was compared with patient derived models of RMC. RMC cell lines (red) cluster with rhabdoid tumor samples (purple). The normal cell line (orange) (CLF_PEDS0005_N) clusters with other normal kidneys profiled in TARGET. **(b)** tSNE analysis of gene-expression array data shows RMC cell lines (red) clustering with RMC patients (blue) and these cluster with other MRT or ATRT samples. **(c)** Glycerol gradients (10-30%) followed by SDS PAGE analysis of rhabdoid tumor cell line, G401, as compared to SMARCB1 wild type cell line HA1E, shows ARID1A is seen in higher fractions when SMARCB1 is expressed. Gradients were performed on patient derived models of RMC with doxycycline-inducible SMARCB1. A similar rightward shift of ARID1A occurs upon re-expression of SMARCB1. These same shifts occur with SMARCA4. These experiments are representative of at least 2 biological replicates. **(d)** Fraction 14 of the glycerol gradients shows a modest increase in SWI/SNF complex members, SMARCC1, SMARCC2, SMARCA4 and ARID1A in HA1E, a SMARCB1 wild type cell line. When SMARCB1 is re-expressed in G401 and RMC lines, a similar pattern is seen. Images are representative of 2 biological replicates. **(e)** Using cell lines with stably transfected inducible SMARCB1, cell viability was assessed with or without expression of SMARCB1 over 8 days. There is no significant difference in SMARCB1 wild type cell line, HA1E. Re-expression of SMARCB1 leads to significant decreases in cell viability as compared to LacZ control in SMARCB1 deficient cancer cell lines G401, CLF_PEDS9001_T, and CLF_PEDS0005. Error bars are standard deviations based on number of samples in parentheses. **(f)** CLF_PEDS0005_T2A cell line show signs of senescence following re-expression of SMARCB1. Images representative of 3 biological replicates.

To determine if the RMC cell lines clustered separately among other SMARCB1-deficient cancers, we performed gene expression analysis of our RMC models and compared them to publicly available datasets of MRT cell lines, patients with RMC, MRT or ATRT or synovial sarcoma (a cancer driven by the fusion oncoprotein SSX-S18 which displaces SMARCB1; **Methods**) (Barretina et al., 2012; Calderaro et al., 2016; Han et al., 2016; Johann et al., 2016; Richer et al., 2017). Using tSNE, we found that the RMC cell lines closely mapped to a French cohort of RMC, MRT and ATRT patients (**Fig. 2B**). These observations demonstrated that RMC cell lines and SMARCB1 deficient patients express similar gene expression programs.

We then assessed the consequences of re-expressing SMARCB1. Specifically, we generated doxycycline-inducible open reading frame (ORF) vectors harboring *SMARCB1* and stably infected our RMC models, G401 (MRT cell line) and HA1E (SMARCB1 wild-type immortalized epithelial kidney cell line) (Hahn et al., 1999). We confirmed that the addition of doxycycline used in our studies did not affect the proliferation of the parental cell lines (**Methods and Supp Fig. 2A-C**).

We used these inducible cell lines to assess the biochemical stability of the SWI/SNF complex by using 10-30% glycerol gradient sedimentation followed by SDS-PAGE (**Fig. 2C-D and Supp Fig. 2D**). In HA1E SMARCB1 wild-type cells, the SWI/SNF complex members SMARCB1, ARID1A and SMARCA4 are robustly expressed at higher molecular weights (e.g. fractions 13-16). In G401 SMARCB1 deficient cells, the SWI/SNF complex is smaller and seen at lower molecular weights (e.g. fractions 10-14). Furthermore, expression of SWI/SNF complex members was modestly decreased in G401, consistent with our prior studies (Nakayama et al., 2017; Wang et al., 2016). In our RMC cell lines, we found that the majority of ARID1A and SMARCA4 was observed in fractions 11-13 similar to what we found in the MRT cell line G401. In addition, we found increased expression and a shift of ARID1A and SMARCA4 to larger fractions 13-15 upon re-expression of SMARCB1 in the RMC lines similar to what we observed in the SMARCB1 wild-type HA1E cell line. We concluded that the composition of the SWI/SNF complex is similar between RMC and other SMARCB1 deficient cancers.

We then used the inducible cell lines to measure the consequence of SMARCB1 re-expression on the viability of the cells. We also generated cell lines with inducible expression of a LacZ control to compare with re-expression of SMARCB1. In the HA1E SMARCB1 wild type cells, we found no significant difference in viability between induction of SMARCB1 versus induction of LacZ using direct counting of viable cells (**Fig. 2E**). In contrast, induction of SMARCB1 in the MRT G401 SMARCB1 deficient cells decreased the number of viable cells by 41%. Similar to G401, we found that re-expression of SMARCB1 in each of the RMC models led to significant decreases in cell viability (37-62%) (**Fig. 2E**). These findings identified that RMC is dependent on the loss of SMARCB1 for survival.

Since MRT cell lines arrest and senesce when SMARCB1 is re-expressed(Betz et al., 2002), we looked for evidence of senescence by staining for senescence-associated acidic β-galactosidase in the RMC cells when SMARCB1 was re-expressed. Following 7 days of SMARCB1 or LacZ re-expression, we stained the cells for β-galactosidase (**Methods**). We failed to observe cells expressing β-galactosidase upon expression of LacZ in the RMC cells, but when SMARCB1 was expressed, we found 44.6% (+/- 17%) of the RMC cells stained for β-galactosidase (**Fig. 2F and Supp Fig. 2E**). These studies showed that re-expression of SMARCB1 in RMC cells leads to senescence.

We then assessed what genes were differentially expressed upon SMARCB1 re-expression in the RMC and MRT cell lines as another way to assess the similarity between these two cancers. We performed RNA-sequencing on the doxycycline-induced SMARCB1 RMC cell lines and compared it to the uninduced cell lines or doxycycline-induced LacZ cell lines. We then re-analyzed our previously published studies of MRT cells with SMARCB1 re-expression and compared it to our RMC cells (Wang et al., 2016). We found 1,719 genes to be significantly different (false discovery rate of <0.25) in the RMC cells and 2,735 genes in MRT cells. We identified 527 genes that significantly overlapped between the RMC and MRT cell lines (hypergeometric p-value less than 4.035e-63; **Supp Data 6**). We then confirmed that these changes in gene expression led to alterations in protein expression by assessing SPARC, p21, c-MYC expression by immunoblotting (**Supp Fig. 2F-G**) (Alpers et al., 2002; Bradshaw and Sage, 2001). These findings confirm that changes in the transcriptome following SMARCB1 re-expression in RMC cell lines are similar to other SMARCB1 deficient cancer cell lines. In sum, these observations indicate that the RMC cell lines are functionally similar to those derived from other SMARCB1 deficient cancers.

### RNAi and CRISPR-Cas9 loss of function screens and small-molecule screens in RMC models identify proteasome inhibition as a vulnerability

MRT, ATRT and RMC are aggressive and incurable cancers. We performed genetic (RNAi and CRISPR-Cas9) and pharmacologic screens to identify druggable targets that would decrease proliferation or survival specifically for these cancers. Specifically, we used the Druggable Cancer Targets (DCT v1.0) libraries and focused on targets that were identified by suppression with RNAi, gene deletion with CRISPR-Cas9 based genome editing, and perturbation by small molecules (Hong et al., 2016; Seashore-Ludlow et al., 2015). We accounted for off-target effects in the RNAi screens by using seed controls for each shRNA.

We performed these three orthogonal screens on both metastatic models of RMC, CLF_PEDS0005_T2A and CLF_PEDS0005_T2B (**Fig. 3A**). We introduced the shRNA DCT v1.0 lentiviral library into these two cell lines and evaluated the abundance of the shRNAs after 26 days using massively parallel sequencing (**Methods**). We confirmed depletion of known common essential genes such as *RPS6* (**Supp Fig. 3A**). We then analyzed the differential abundance between the experimental and seed control shRNAs to collapse individual shRNAs to consensus gene dependencies with RNAi Gene Enrichment Ranking (RIGER) (Luo et al., 2008). Of 444 evaluable genes, 72 genes scored with a RIGER p-value <0.05 in CLF_PEDS0005_T2A and 74 genes scored in CLF_PEDS0005_T2B.

**Fig. 3:**
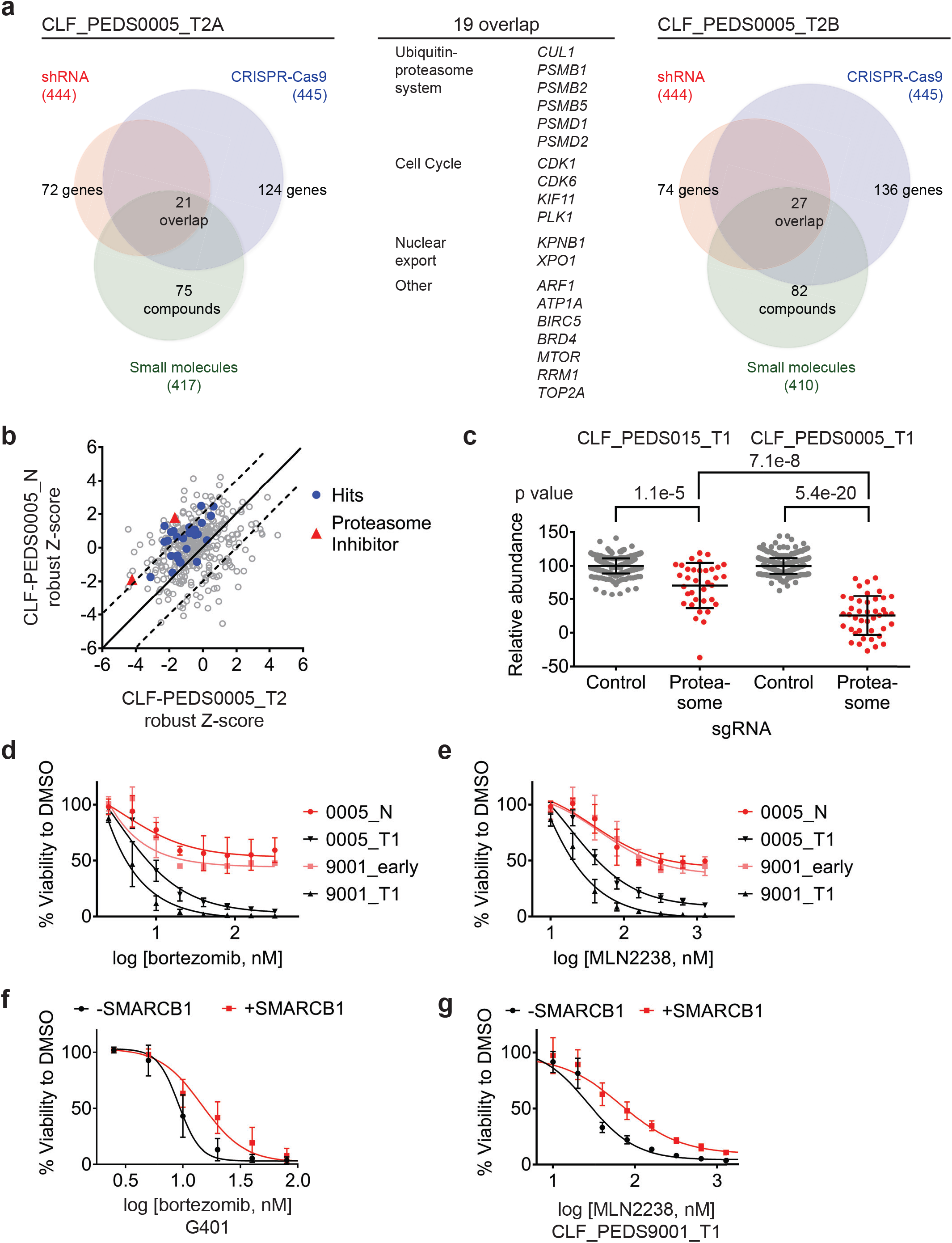
Functional genomic screens identify inhibition of the proteasome as a vulnerability in RMC. **(a)** Left: RNAi suppression of 444 evaluable genes (red) identifies 72 genes that when suppressed caused a significant viability loss in CLF_PEDS0005_T2A. Genomic indels created by CRISPR-Cas9 in 445 evaluable genes (blue) identify 124 genes that cause a viability loss. RNAi and CRISPR-Cas9 screens were performed in biological replicates. Small-molecule screen (performed in technical replicates) with 417 evaluable compounds (green) identifies 75 compounds that lead to significant viability loss. Twenty one genes overlap across these three screens. Right: The same screens were performed with CLF_PEDS0005_T2B and 27 genes were found to be significantly depleted when suppressed by RNAi, genomically deleted by CRISPR-Cas9 or inhibited when treated with a small molecule. Center: List of 19 genes that overlapped between functional screens from CLF_PEDS0005_T2A and CLF_PEDS0005_T2B can be categorized into genes involving the ubiquitin-proteasome system, cell cycle and nuclear export. **(b)** Comparison of Z-score normalized small-molecule screens between CLF_PEDS0005_T2 and CLF_PEDS0005_N (normal isogenic cell line). Hits (blue) and proteasome inhibitors (red). Each dot is representative of the average of two technical replicates. **(c)** Relative log2 fold change in abundance from CRISPR-Cas9 screens between sgRNA controls (grey) and genes in the DCT v1.0 screen involving the proteasome (red). Data is taken at 23 days following selection and compared to an early time point. As compared to the undifferentiated sarcoma cell line CLF_PEDS015T, inhibition of the proteasome subunits leads to a more profound viability loss as compared with controls. Each dot is representative of a minimum of 2 biological replicates. **(d)** Short term cultures of the normal cell line (CLF_PEDS0005_N) or early passage of the heterogenous cell line (CLF_PEDS9001_early) were compared for assessment of viability to the primary tumor cell lines following treatment with bortezomib. Two-tailed t-test p-value = 0.008 for PEDS0005 and two-tailed t-test p-value = 4.76e-5 for PEDS9001. Error bars represent standard deviations from two biological replicates. **(e)** Short term cultures of the normal cell line (CLF_PEDS0005_N) or early passage of the heterogenous cell line (CLF_PEDS9001_early) were compared for assessment of viability to the primary tumor cell lines following treatment with MLN2238. Error bars represent standard deviations from two biological replicates. **(f)** Re-expression of SMARCB1 in G401 leads to a rightward shift in the dose-response curve with bortezomib compared with uninduced cells. Error bars represent standard deviations from two biological replicates. **(g)** Re-expression of SMARCB1 in CLF_PEDS9001_T leads to a rightward shift in the dose-response curve with MLN2238 compared with uninduced cells. Error bars represent standard deviations from three biological replicates.

In parallel, we introduced the CRISPR-Cas9 DCT v1.0 lentiviral library to determine the differential representation of the CRISPR-Cas9 sgRNAs between 6 and 23 days to identify genes depleted or enriched in this screen by massively parallel sequencing (**Methods**). We confirmed that the distribution of sgRNAs among biological replicates was highly correlated (**Supp Fig. 3B-E**). When compared to the controls, there was significant depletion of essential genes such as *RPS6* (**Supp Fig. 3F-G**). We used RIGER to collapse the individual sgRNAs to consensus gene dependencies and found 124 genes (of a total of 445 evaluable genes) and 136 genes (of a total of 445 evaluable genes) with a RIGER p-value <0.05 in CLF_PEDS0005_T2A and CLF_PEDS0005_T2B cell lines, respectively.

We performed a small-molecule screen using a library of 440 compounds that have known targets in the RMC cells (CLF_PEDS0005_T2A and CLF_PEDS0005_T2B) (Hong et al., 2016). This library includes 72 FDA approved compounds, 100 compounds in clinical trials and 268 probes based on our prior studies (Hong et al., 2016). We calculated an area under the curve (AUC) based on an 8-point concentration range and considered AUCs <0.5 as significant. Of the evaluable compounds, 75 (18%) compounds significantly decreased cell viability in CLF_PEDS0005_T2A and 82 (20%) compounds significantly decreased cell viability in CLF_PEDS0005_T2B.

We then looked for genes or targets of the small molecules that scored in all three of the RNAi, CRISPR-Cas9 and small-molecule screens. We identified 21 genes in CLF_PEDS0005_T2A and 27 genes in CLF_PEDS0005_T2B (**Data File S7**) of which 19 genes scored in both screens (**Fig. 3A**). Among the 19 genes were components of the ubiquitin-proteasome system (e.g. *PSMB1, PSMB2, PSMB5, PSMD1, PSMD2*, and *CUL1*), regulators of the cell cycle (*CDK1, CDK6, KIF11* and *PLK1*), genes involved in nuclear export (*KPNB1* and *XPO1*) and *ARF1, ATP1A, BIRC5, BRD4, MTOR, RRM1* and *TOP2A.*

To eliminate small molecules and targets that affect normal renal tissue, we screened the normal cell line (CLF_PEDS0005_N) with the small-molecule library. We calculated the robust Z-scores for these screens in relationship to the Cancer Cell Line Encyclopedia (CCLE) to normalize the responses to various compounds (Barretina et al., 2012; Rees et al., 2016; Seashore-Ludlow et al., 2015). We then compared the results of this small-molecule screen with the RMC cancer cell lines (CLF_PEDS0005_T2A and CLF_PEDS0005_T2B). We found that the tumor cells were differentially sensitive (up to two standard deviations) upon treatment with proteasome inhibitors, bortezomib and MLN2238, when compared to the normal cell line (**Fig. 3B**). These findings suggest that the vulnerability to proteasome inhibition may be dependent on loss of SMARCB1.

### Validation of proteasome inhibition as a specific dependency in SMARCB1 deficient cancers

To validate the dependency of SMARCB1 deficient tumors to the ubiquitin-proteasome system, we assessed the consequences of inhibiting proteasome function on survival of the primary tumor cell line, CLF_PEDS0005_T1, by deleting components of the proteasome with CRISPR-Cas9. We compared these findings with a model of undifferentiated sarcoma, CLF_PEDS015T, that does not harbor mutations in *SMARCB1* (Hong et al., 2016). We scaled the results based on the non-targeting sgRNA negative controls and *RPS6* as a common essential gene (Hart et al., 2015). Compared to the control sgRNAs, there was an average decrease of 29% in viability in CLF_PEDS015T while there was an average decrease of 74% in CLF_PEDS0005_T1 (**Fig. 3C; Methods**). Although deletion of the proteasome members affected proliferation in all of the models (two tailed t-test p = 1.1e-5 for CLF_PEDS015T and p = 5.4e-20 for CLF_PEDS0005_T1), we found that suppression of proteasome components affected the RMC model CLF_PEDS0005_T1 to a statistically greater degree (two tailed t-test p = 7.1e-8). We subsequently validated that gene deletion by CRISPR-Cas9 of *PSMB5*, one of the primary targets of proteasome inhibitors, in the CLF_PEDS0005_T2A and CLF_PEDS9001_T1 cell lines led to decreased viability (**Supp Fig. 3H-I**).

We then determined whether this vulnerability to proteasome inhibition was specific to the loss of SMARCB1. We treated the normal cell line, CLF_PEDS0005_N, and an early passage of CLF_PEDS9001_T1 while it was a heterogenous population and retained *SMARCB1* (**Supp Fig. 3J**) with bortezomib or the second-generation proteasome inhibitor, MLN2238. We observed significantly decreased sensitivity to the proteasome inhibitors in the *SMARCB1* retained isogenic cell lines as compared to the *SMARCB1* deficient cell lines (**Fig. 3D-E**). We then treated our SMARCB1-inducible RMC and MRT cell lines with DMSO, bortezomib or MLN2238. Re-expression of SMARCB1 led to a decrease in sensitivity to bortezomib or MLN2238 as compared to the isogenic SMARCB1 deficient lines (**Fig. 3F-G**). The observed differential resistance to SMARCB1 re-expression was between 2-3-fold with either bortezomib or MLN2238 (**Supp Fig. 4A-B**). We concluded that re-expression of SMARCB1 partially rescued the sensitivity of MRT or RMC cell lines to proteasome inhibition.

We then compared the results of small-molecule screens performed in SMARCB1-deficient cancer cell lines in CCLE to the rest of the CCLE cell lines (n=835). We found that SMARCB1-deficient cell lines were significantly more sensitive (two-tailed t-test p-value = 0.011) to treatment with MLN2238 than non-multiple myeloma CCLE cell lines (**Fig. 4A**) (Rees et al., 2016; Seashore-Ludlow et al., 2015). The degree of sensitivity was similar to that of multiple myeloma cell lines which are known to be sensitive to proteasome inhibition (Dimopoulos et al., 2016). These findings confirm that SMARCB1 deficient cell lines are selectively vulnerable to proteasome inhibition.

**Fig. 4:**
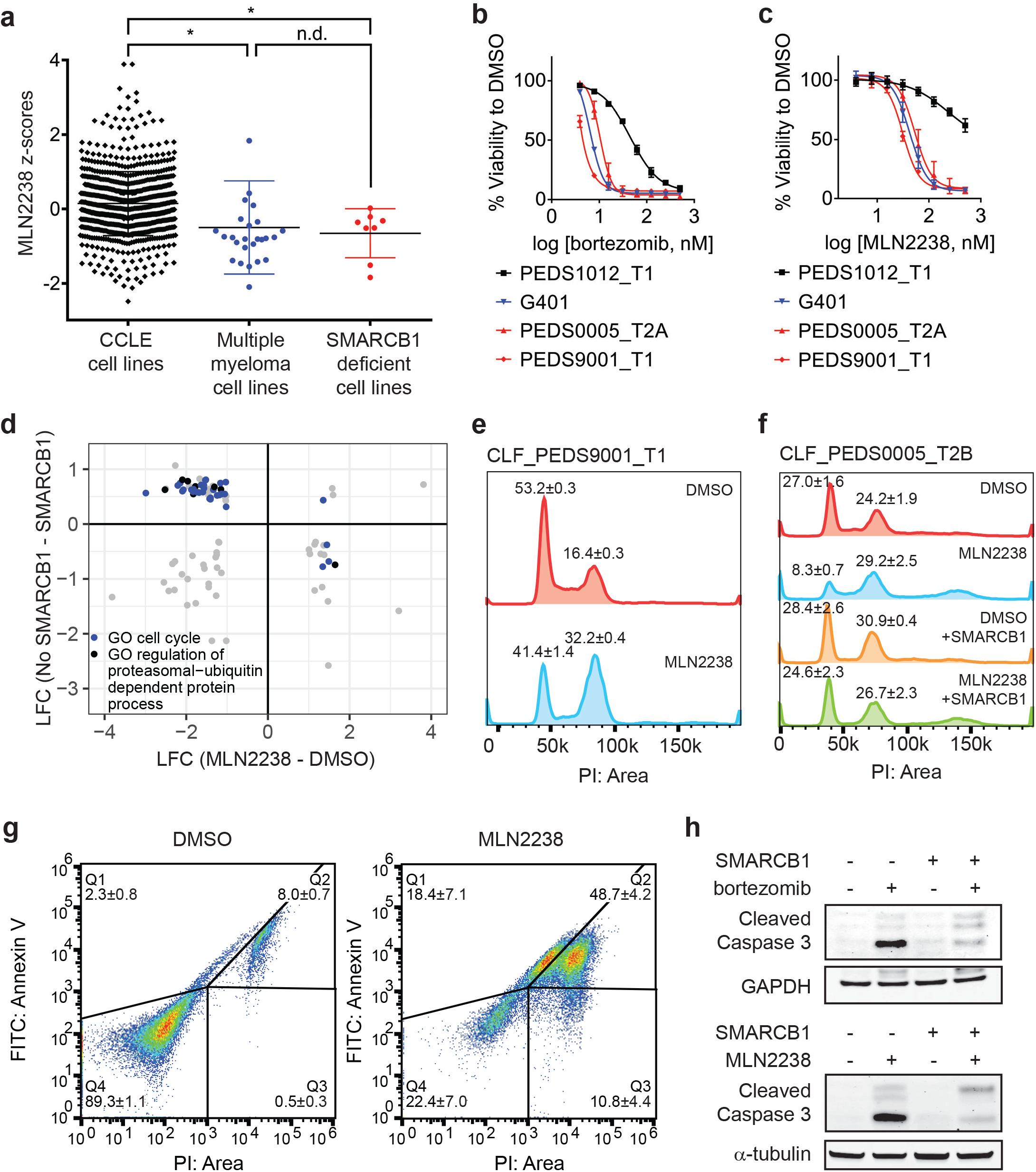
Proteasome inhibitors such as MLN2238 are specific to SMARCB1 deficient cancers and lead to cell cycle arrest and programmed cell death. **(a)** Multiple myeloma cell lines and SMARCB1 deficient lines are similarly sensitive to proteasome inhibitor, MLN2238. Both are significantly different from other CCLE cell lines based on Z-score normalized sensitivity. * two-tailed t-test p-value < 0.05. n.d. no difference. **(b)** Early passage Wilms tumor (SMARCB1 wild type) cell line CLF_PEDS1012_T1 is not as sensitive to treatment with bortezomib compared with RMC and MRT cell lines. Error bars represent standard deviations following at least two biological replicates. **(c)** Early passage Wilms tumor (SMARCB1 wild type) cell line CLF_PEDS1012_T1 is not as sensitive to treatment with MLN2238 compared with RMC and MRT cell lines. Error bars represent standard deviations following at least two biological replicates. **(d)** Analysis of differential genes when SMARCB1 was re-expressed in SMARCB1 deficient cancers compared with differential genes when SMARCB1 deficient cancers were treated with MLN2238. Gene sets enriched based on GO based GSEA involved the cell cycle (blue) and regulation of the ubiquitin-proteasome system (black). **(e)** Treatment with proteasome inhibitor, MLN2238 for 24 hours leads to G2/M arrest in CLF_PEDS9001_T1. Values shown represent the percent of cells in G1 or G2/M. Error values shown are standard deviations from two biological replicates. **(f)** Treatment with MLN2238 for 24 hours leads to G2/M arrest in CLF_PEDS0005_T2B which can be prevented by re-expression of SMARCB1. Error values shown are standard deviations from two biological replicates. **(g)** Treatment of CLF_PEDS9001_T1 with MLN2238 for 48 hours leads to increased frequency of cells with Annexin V / PI staining and PI only staining. Error values shown are standard deviations from two biological replicates. **(h)** G401 cells stably infected with inducible SMARCB1 treated with either bortezomib or MLN2238 induce cleaved caspase 3 compared with DMSO controls after 24 hours. When SMARCB1 is re-expressed, cleaved caspase 3 levels are decreased compared to uninduced cell lines. Blots are representative of a minimum of 2 biological replicates.

We then performed *in vitro* studies to confirm the findings from these high throughput small molecule screens. We treated an additional 6 SMARCB1 deficient cell lines (4 MRT and 2 ATRT) with bortezomib. We compared these results to H2172, a lung cancer cell line that was not sensitive to proteasome inhibition in CCLE small-molecule screens, and RPMI8226, an established multiple myeloma cell line that is responsive to proteasome inhibition (Hideshima et al., 2001). We found that our SMARCB1 deficient cell lines exhibited single digit nanomolar sensitivity to proteasome inhibition similar to that observed in the multiple myeloma cell line RPMI8226 (**Supp Fig. 4C**). In contrast, we found that the IC50 in H2172 was at least 3-fold higher. Since there are no SMARCB1 wildtype pediatric kidney cancer cell lines in CCLE, we compared the sensitivity to bortezomib or MLN2238 in our RMC models with wildtype SMARCB1 patient-derived Wilms tumor cell line, CLF_PEDS1012_T (**Fig. 4B-C and Supp Fig. 3J**). We found that CLF_PEDS1012_T was more resistant to proteasome inhibition as compared to our RMC models and MRT cell line, G401. These findings suggest that SMARCB1-deficient cells are more sensitive to proteasome inhibition.

### Proteasome inhibition leads to cell cycle arrest in G2/M and subsequent cell death

We then studied how SMARCB1 loss leads to a dependency on the ubiquitin-proteasome system. Since activation of c-MYC has been observed in SMARCB1-deficient cancers (Cheng et al., 1999; Genovese et al., 2017), we assessed how c-MYC levels are altered upon proteasome inhibition. Compared to RPMI8226, a multiple myeloma cell line that relies on c-MYC for survival (Tagde et al., 2016), we failed to observe suppression of c-MYC protein levels following bortezomib or MLN2238 treatment (**Supp Fig. 4D**). We then assessed c-MYC expression levels by RNA-sequencing in the G401, CLF_PEDS9001T, and CLF_PEDS00005_T1 cell lines following treatment with MLN2238 and found that c-MYC levels were increased after MLN2238 treatment (**Supp Fig. 4E**). In our models of RMC and MRT, these findings suggest proteasome inhibition does not lead to suppression of c-MYC.

We then assessed if the SWI/SNF complex is altered upon treatment with a proteasome inhibitor. We treated uninduced and induced SMARCB1 cells with DMSO or a proteasome inhibitor. We failed to see a consistent significant change in total or nuclear protein when immunoblotting for SWI/SNF complex members (BAF57, BAF60A, BAF155, BAF170, ARID1A and SMARCA4) other than increases in SMARCB1 levels upon doxycycline treatment (**Supp Fig. 5A-B**). These findings suggest that inhibition of the proteasome in SMARCB1 deficient cancers and its subsequent resistance upon SMARCB1 expression does not alter the total or nuclear levels of the SWI/SNF complex.

ER stress has been implicated as a mechanism by which proteasome inhibitors act on multiple myeloma cells (Obeng et al., 2006). We saw an increase in protein expression of markers of ER stress, GRP78 and IRE1α, following treatment with MLN2238 (**Supp Fig. 5C**). Upon re-expression of SMARCB1 and subsequent treatment with a proteasome inhibitor, we did not see changes in either GRP78 or IRE1α protein levels (**Supp Fig. 5C**). These observations suggest that although ER stress markers are elevated upon proteasome inhibition in SMARCB1-deficient cell lines, they are not rescued by SMARCB1 re-expression.

We performed gene ontology (GO) based Gene Set Enrichment Analysis (GSEA)(Subramanian et al., 2005) on the 527 significantly altered genes upon re-expression of SMARCB1 (**Supp Data 6**) to identify classes of gene function enriched in this group of genes. We identified numerous gene sets that involved the ubiquitin-proteasome system (**Supp Data 8**). We then performed RNA-sequencing on G401 and the RMC cell lines treated with DMSO or MLN2238. We identified 1,758 genes which were significantly (FDR<0.1) up- or down-regulated upon treatment with MLN2238 (**Supp Data 9**). We compared the 527 differentially expressed genes identified upon re-expression of SMARCB1 with the 1,758 differentially expressed genes identified upon treatment with MLN2238 and identified 92 genes which overlapped. Of these genes, we identified 63 genes which were differentially expressed with re-expression of SMARCB1 and were differentially expressed with treatment with MLN2238 (**Fig. 4D**). From this refined gene set, we performed GO-based GSEA (Subramanian et al., 2005) and found significant enrichment (adjusted p-value ranging from 0 to 0.0061) in gene sets involving the cell cycle (**Supp Data 10**).

We then assessed the cell cycle by DNA content with cells treated with DMSO or MLN2238 for 24 hours as proteasome inhibitors have been found to cause a G2/M cell cycle arrest in lymphomas, colorectal carcinomas, hepatocellular carcinomas, and glioblastoma multiforme (Augello et al., 2018; Bavi et al., 2011; Gu et al., 2017; Yin et al., 2005). We observed a significant shift in cells to G2/M (two-tailed t-test p-value 0.0005; **Fig. 4E**). Upon re-expression of SMARCB1, we saw that this phenotype was rescued (**Fig. 4F**). By 48 hours, we saw a significant increase in markers of programed cell death such as Annexin V and PI positive cells (**Fig. 4G**).

We found that treatment with a proteasome inhibitor led to increased cleaved caspase-3 levels in addition to changes to Annexin V and PI, suggesting that inhibition of the ubiquitin-proteasome system leads to programmed cell death (**Fig. 4H**). We then asked whether restoration of SMARCB1 expression inhibited cleaved-caspase 3 activation. We found that induction of cleaved caspase-3 was less pronounced when SMARCB1 was re-expressed (**Fig. 4H**). These observations suggest that proteasome inhibitors initially lead to a SMARCB1-dependent G2/M cell cycle arrest and subsequent programmed cell death.

### Proteasome inhibitor induced G2/M arrest is mediated in part by inappropriate cyclin B1 degradation driven by a dependency on UBE2C

We subsequently searched for genes related to the ubiquitin-proteasome system and SMARCB1 function. We defined a set of 204 genes that were upregulated when comparing the log2 fold change between SMARCB1 deficient cells to SMARCB1 re-expressed cells in RMC and MRT cell lines (**Supp Data 6**). We then took this set of 204 genes and examined the Project Achilles (genome scale CRISPR-Cas9 loss of function screens) DepMap Public 18Q3 dataset (Meyers et al., 2017) to determine whether any SMARCB1 deficient cell lines required expression of these genes for survival. This dataset included loss of function screens from 485 cancer cell lines and included three ATRT SMARCB1-deficient cancer cell lines: COGAR359, CHLA06ATRT and CHLA266. We found that SMARCB1 deficient cancer cell lines required *UBE2C*, a ubiquitin-conjugating enzyme, for survival. We noted that these cell lines were in the top 5% of cell lines (n=485) that required *UBE2C* for survival (empirical Bayes moderated t-test p-value = 0.00016; **Fig. 5A-B**). These observations suggested that cancer cell lines that lack SMARCB1 were also dependent on UBE2C.

**Fig. 5:**
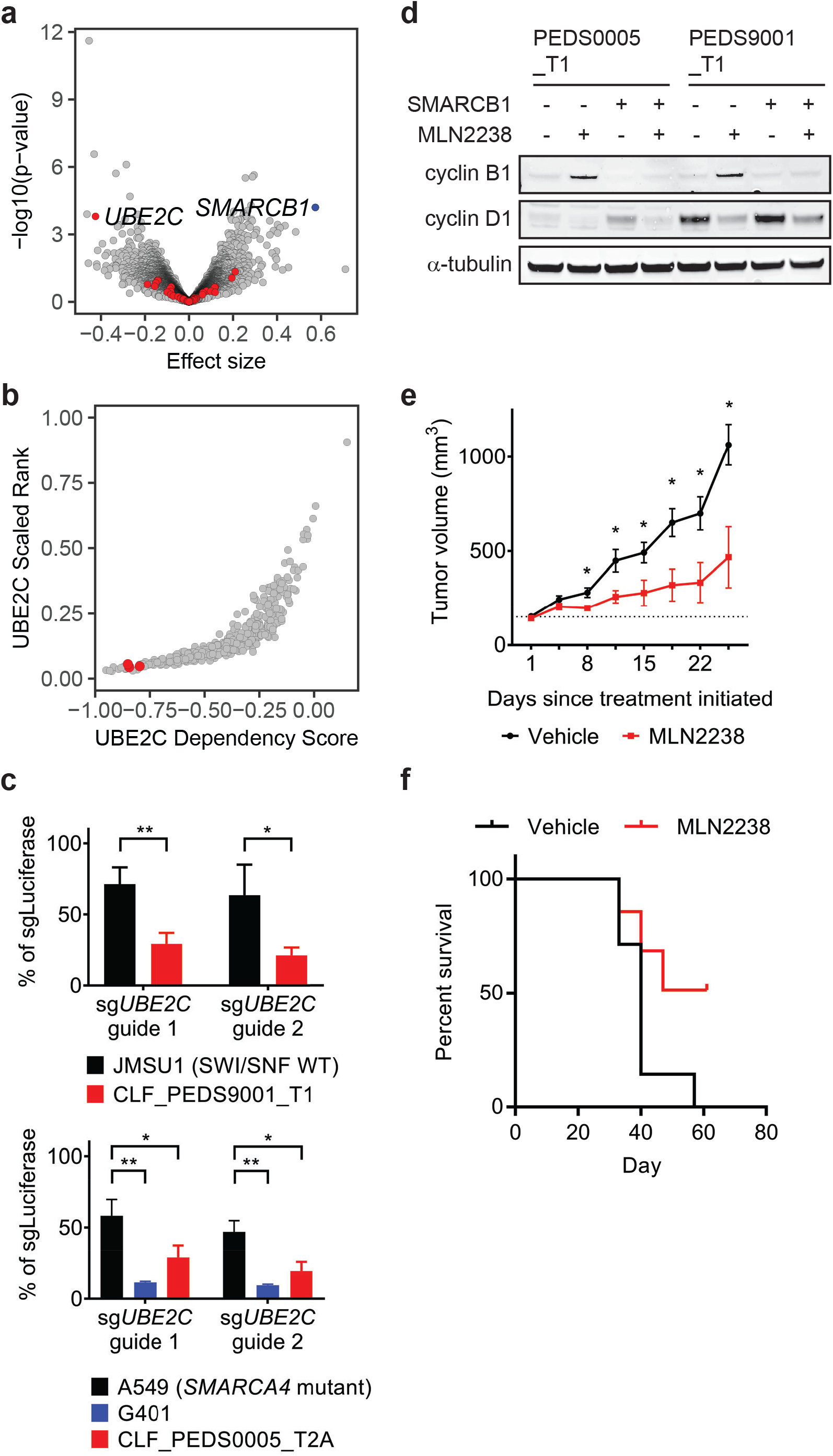
Cell cycle arrest secondary to proteasome inhibition acts via UBE2C/cyclin B1 dysregulation and proteasome inhibition with MLN2238 is effective *in vivo.* **(a)** 204 genes were significantly upregulated when comparing the log2 fold change between SMARCB1 deficient cells to SMARCB1 re-expressed cells (**Supp Data 6**). We assessed how loss of these genes affected viability in 3 cell lines with loss of SMARCB1 as compared to the rest of the 482 cell lines in Project Achilles DepMap 18Q3, a genome-wide loss of function screen using CRISPR-Cas9 and calculating an effect size (e.g. differential of the 3 cell lines to 482 cell lines). A negative effect size identifies genes when deleted are required for cells for survival and the 204 genes are identified in red. Deletion of *UBE2C* was significantly depleted. Deletion of *SMARCB1* serves as a positive control in these SMARCB1 deficient cancers as these cell lines have loss of SMARCB1. **(b)** SMARCB1 deficient lines are in the top 5% of cell lines ranked by how dependent they are on *UBE2C* based on Project Achilles DepMap 18Q3 dataset. Three ATRT cancer cell lines (red dots; CHLA266, CHLA06, COGAR359) were compared to 485 cell lines profiled in Project Achilles (CERES dataset 18Q3). **(c)** Gene deletion of *UBE2C* by CRISPR-Cas9 leads to significant viability defects in RMC and MRT cell lines as compared to either SWI/SNF wt cell line, JMSU1 (day 10), or *SMARCA2* mutant cell line, A549 (day 6). Error bars shown are standard deviations from two biological replicates. * indicates a two-tailed t-test p-value < 0.05 and ** <0.005. **(d)** Treatment with proteasome inhibitor, MLN2238, leads to upregulation of cyclin B1 and this phenotype is rescued upon SMARCB1 re-expression in both CLF_PEDS0005_T1 and CLF_PEDS9001_T1. Cyclin D1 is included as a control to ensure that the effects of the proteasome are specific to cyclin B1. Blots are representative of two biological replicates. **(e)** G401 xenograft tumor growth over time shows that treatment effects from MLN2238 can be seen as early as 8 days from treatment initiation as compared to vehicle control. Dotted line reflects average volume of 148mm^3^. Error bars are reflective of standard error of the mean. * indicates Mantel-Cox p-value <0.05. **(f)** Kaplan-Meier curves from mice with G401 xenograft tumors treated with either vehicle or MLN2238.

Since the 3 cell lines profiled were ATRTs, we validated that the RMC cell lines were also dependent on *UBE2C* for survival. We generated sgRNAs specific for *UBE2C* and assessed viability by cell counting following gene deletion. We saw a significant decrease in cell viability in SMARCB1 deficient cell lines as compared to urothelial carcinoma cell line, JMSU1 (SWI/SNF wild type), or non-small cell lung cancer cell line, A549 *(SMARCA4* mutant) (two-tailed t-test p-values 4.5e-5 and 4.6e-5; **Fig. 5C and Supp Fig. 6A**).

UBE2C serves as the E2 enzyme which adds the first ubiquitin (Ub) to cyclin B1 for degradation (Dimova et al., 2012; Grice et al., 2015). Cyclin B1 degradation is required in G2/M at the end of metaphase to enter anaphase(Chang et al., 2003). Our integrated RNAi, CRISPR-Cas9 and small molecule screens identified that our RMC models required expression of *PLK1* and *CDK1*, genes involved in G2/M, for survival (**Fig. 3A-B**), and prior studies have identified that inhibition of *PLK1* in ATRT or MRT cells lead to arrest in G2/M (Alimova et al., 2017; Morozov et al., 2007). Treatment of the RMC cell lines with MLN2238 led to accumulation of cyclin B1 as compared to cyclin D1 suggesting that MLN2238 inhibits degradation of cyclin B1 (**Fig. 5D**). When we re-expressed SMARCB1, we found that cyclin B1 levels were unchanged upon MLN2238 treatment. Although APC/C serves as the E3 ligase for cyclin B1, genetic deletion of APC/C in Project Achilles showed that APC/C was an essential gene across all cancer cell lines. These findings suggest SMARCB1 deficient cancer cells require UBE2C expression for survival, in part by regulating cyclin B1 stability.

### Effects of proteasome inhibition in vivo

These studies identify a lethal interaction between suppressing the ubiquitin-proteasome system and SMARCB1-deficient cancers *in vitro.* The doses used in this study were based on *in vitro* studies of multiple myeloma or lymphoma cell lines (Chauhan et al., 2011; Garcia et al., 2016; Hideshima et al., 2003, 2001). For patients with primary or refractory multiple myeloma, use of proteasome inhibitors has led to significant clinical responses (Jagannath et al., 2004; Moreau et al., 2016; Richardson et al., 2005, 2003). We reasoned that if our SMARCB1 deficient cancers were susceptible to proteasome inhibitors at similar *in vitro* dosing, we would also see similar *in vivo* responses. We first determined whether these doses led to proteasome inhibition by assessing the ability of these cell lines to cleave Suc-LLVY-aminoluciferin. We found that treatment of the SMARCB1 deficient cell lines with either bortezomib or MLN2238 led to inhibition of the proteasome to a similar extent observed when the multiple myeloma cell line RPMI8226 was treated (**Supp Fig. 6B; Methods**). We also simulated the pharmacodynamics of proteasome inhibitors *in vivo* by treating cells *in vitro* with a pulse dose of proteasome inhibitors as has been performed in multiple myeloma and chronic myeloid leukemia cell lines (Crawford et al., 2014; Kuhn et al., 2007; Shabaneh et al., 2013). We found that upon treatment with MLN2238, SMARCB1 deficient cells arrested in G2/M, which led to cell death as measured by Annexin V/PI staining and led to accumulation of cyclin B1 similar to treatment with a continuous dose of MLN2238 (Methods; **Supp Fig. 6C-F and 7A-C**).

We subsequently performed *in vivo* studies to confirm the effect of monotherapy with proteasome inhibition in tumor xenografts. We used the rhabdoid tumor cell line, G401, for our *in vivo* studies because we noted that the primary tumor cell lines (CLF_PEDS0005_T1 and CLF_PEDS9001_T) did not form subcutaneous xenograft tumors in immunodeficient mice (**Supp Fig. 7D-E**). We allowed tumors to achieve an average volume of 148 mm^3^ treated mice with either vehicle or MLN2238 (7 mg/kg twice a week) and measured tumors twice weekly. Treatment with MLN2238 induced significant tumor stabilization (n=4) or regression (n=3) when assessed either by relative growth (two-tailed t-test p-value 0.038; **Supp Fig. 7F**) or by absolute tumor volume (two-tailed t-test p-value 0.012; **Fig. 5E**) and did not induce significant changes in body weight as compared to vehicle controls (two-tailed t-test p-value 0.154; **Supp Fig. 7G**). Furthermore, there was a significant survival for mice treated with MLN2238 (p-value 0.0489 by log-rank (Mantel-Cox) test; Fig. 5F). Combined, these results demonstrate that MLN2238 induces tumor regression in SMARCB1 deficient tumors *in vivo.*

## Discussion

We have developed faithful patient-derived models of RMC which have been genomically validated using PCR-free WGS, WES, RNA-sequencing and gene expression profiling. We have shown that these models are dependent upon the loss of SMARCB1 for survival. Re-expression of SMARCB1 in RMC leads to a significant decrease in cell counts and a senescence phenotype. Biochemically, re-expression of SMARCB1 in RMC leads to stabilization of the SWI/SNF complex in the same manner as for MRT. Diagnostically, patients with RMC are often misdiagnosed with renal cell carcinoma (RCC) due to the rarity of RMC, the lack of access to SMARCB1 histological stains and unknown sickle cell status (Beckermann et al., 2017). Although SMARCB1 is currently included in targeted sequencing efforts nationwide(“AACR Project GENIE: Powering Precision Medicine through an International Consortium,” 2017), our studies along with prior studies (Calderaro et al., 2016; Carlo et al., 2017) suggest that conventional target exome sequencing may fail to identify patients with RMC.

Patients with RMC and other SMARCB1 deficient cancers have a poor prognosis despite aggressive multi-modal therapy. Using genetic and pharmacologic screens in these RMC models, we identified the ubiquitin-proteasome system as a specific vulnerability in RMC. When we looked more broadly at other SMARCB1 deficient cancers such as MRT and ATRT, we found that these models were similarly sensitive to inhibition of the ubiquitin-proteasome system. Re-expression of SMARCB1 partially rescued the sensitivity to proteasome inhibitors in RMC and MRT models.

Prior studies have tested bortezomib in the G401 MRT cell line in an *in vitro* setting or implicated MYC signaling and downstream activation of ER stress as a mechanism for sensitivity to proteasome inhibitors in Kras/Tp53 mutant pancreatic cancers with Smarcb1 deficiency (Genovese et al., 2017; Moreno and Kerl, 2016). However, the background of mutant KRAS may be contributing to these findings as mutant HRAS or KRAS cancers are sensitive to enhanced proteotoxic stress and ER stress (De Raedt et al., 2011; Denoyelle et al., 2006). Furthermore, KRAS mutant cancers depend on several proteasome components in genome scale RNAi screens (Aguirre and Hahn, 2018; Barbie et al., 2009; Luo et al., 2009). Our studies have identified that the ubiquitin-proteasome system is a core vulnerability among a compendium of druggable targets as tested by orthogonal methods of RNA interference, CRISPR-Cas9 gene deletion or small molecule inhibition. We found that proteasome inhibition in SMARCB1-deficient cancer cell lines result in cells accumulating in G2/M due to inappropriate degradation of cyclin B1.

Taken together, these results explain the sensitivity of SMARCB1 deficient lines to proteasome inhibitors such as MLN2238 *in vitro* and *in vivo.* There have been case reports of one adult and 2 children with RMC who exhibited extraordinary responses of 2-7 years following diagnosis after empiric therapy with bortezomib has been used either as monotherapy or in combination with chemotherapy (Carden et al., 2017; Ronnen et al., 2006). Our results suggest that future clinical trials could incorporate oral proteasome inhibitor such as MLN2238 for patients with RMC and potentially more broadly across SMARCB1 deficient cancers. Finally, our studies show how interaction with patient advocacy foundations and online direct-to-patient consent process may serve as a model to study other rare cancers and lead to advances in rare tumor research (Sharifnia et al., 2017).

## Materials and Methods

### Derivation of RMC models

Patients assented or families consented to IRB approved protocols. Patient PEDS0005 whole blood, adjacent normal kidney and tumor tissue were obtained within 6 hours as part of the nephrectomy following neoadjuvant chemotherapy. Upon relapse, pleural fluid was obtained from a palliative thoracentesis. The tumor and adjacent normal kidney tissue was minced into 2-3mm^3^ cubes. CLF_PEDS0005 primary tissue was then dissociated as previously described and cultured in both RPMI media containing 10% FBS (Sigma) or DMEM/F-12 media containing ROCK inhibitor, Y-27632, insulin, cholera toxin, 5% FBS, and penicillin/streptomycin (Liu et al., 2012). Samples were gently passaged when cultures achieved 80-90% confluence. The normal kidney cell line was named CLF_PEDS0005_N. The tumor kidney cell line was named CLF_PEDS0005_T1. For the pleural fluid sample, samples were grown initially in conditioned media as previously published (Liu et al., 2012). Adherent and suspension cells were continuously passaged when cells reached confluence. By passage 5, cells were noted to be growing as adherent and suspension cells, and these were sub-cultured to yield CLF_PEDS0005_T2A and CLF_PEDS0005_T2B, respectively. Samples were then transitioned to DMEM/F-12 media or RPMI as above at passage 13.

Patient PEDS9001 whole blood, adjacent normal kidney and tumor tissue were obtained similarly to PEDS0005. Discarded tissue from the nephrectomy following neoadjuvant chemotherapy was sent to our institution within 24 hours of resection. For the normal kidney and tumor tissue, samples were minced to 2-3mm^3^ and plated onto 6 well plates (Corning, NY). Following mincing, tumor samples were cultured without further digestion. Tumor samples were grown in culture in the DMEM/F-12 media as described above to yield CLF_PEDS9001_T1, while the normal samples yielded a cell culture that matched the tumor cell line.

### Breakapart fluorescent *in situ* hybridization (FISH)

We developed a custom dual-color breakapart FISH probe, using BAC probes surrounding SMARCB1 at 22q11.23: RP11-662F7 (telomeric to SMARCB1, labeled in green) and RP11-1112A23 (centromeric to SMARCB1, labeled in red) (Empire Genomics, Buffalo, NY). The probe set was hybridized to a normal control to confirm chromosomal locations and to determine the frequency of expected fusion signals in normal cells. 50 nuclei were scored by two independent observers (n=100 per cell line) in the CLF_PEDS0005 and CLF_PEDS9001 models.

### Whole Exome Sequencing

We performed whole exome sequencing (WES) from genomic DNA extracted from whole blood, normal / tumor tissues, and our patient derived cell lines as noted in the text. One ug of gDNA (as measured on a Nanodrop 1000 (Thermo Fisher Scientific)) was used to perform standard (~60x mean target coverage for normal) or deep (~150x mean target coverage for tumor and cell lines) WES. Illumina (Dedham, MA) chemistry used.

### Whole Genome Sequencing

We performed PCR-free whole genome sequencing (WGS) from gDNA extracted from whole blood, normal / tumor tissues, and our patient derived cell lines as noted in the text. Two ug of gDNA was used to perform standard (for normal) or deep (for tumor and cell lines) coverage. Illumina (Dedham, MA) HiSeq X Ten v2 chemistry was used. Average coverage is indicated in the text.

### RNA-sequencing

For **Fig. 2A**, samples were processed using Illumina TruSeq strand specific sequencing. We performed poly-A selection of mRNA transcripts and obtained a sequencing depth of at least 50 million aligned reads per sample. For SMARCB1 re-expression RNAseq experiments and MLN2238 vs DMSO treated experiments, samples were collected as biological replicates or triplicates. RNA was extracted using Qiagen RNeasy Plus Mini Kit (Qiagen, Hilden, Germany). RNA was normalized using the Qubit RNA HS Assay (Thermo Fisher Scientific). Five hundred ng of normalized RNA was subsequently used to create libraries with the Kapa Stranded mRNA-seq kit (Kapa Biosystems, KK8420; Wilmington, MA). cDNA libraries were then quantitatively and qualitatively assessed on a BioAnalyzer 2100 (Agilent, Santa Clara, CA) and by qRT-PCR with Kapa Library Quantification Kit. Libraries were subsequently loaded on an Illumina HiSeq 2500 and achieved an average read depth of 10 million reads per replicate.

### Genomic analyses

WGS - Samples were aligned to Hg19. SvABA v0.2.1 was used to identify large deletions and structural variations (Forbes et al., 2015). WES – Samples were aligned to Hg19. Samples were analyzed using GATK v4.0.4.0 for copy number variation (CNV), single nucleotide polymorphism (SNP) and indel identification across our RMC samples simultaneously using filtering parameters set by GATK (Broad Institute, Cambridge, MA) (McKenna et al., 2010). MuTect2 was used to identify candidate somatic mutations and these were correlated with COSMIC (Forbes et al., 2015). RNA – CLF_PEDS0005 and CLF_PEDS9001 samples and TARGET Wilms and Rhabdoid tumor samples (dbGaP phs000218.v19.p7) were aligned or re-aligned with STAR and transcript quantification performed with RSEM. The TARGET initiative is managed by the NCI and information can be found at http://ocg.cancer.gov/programs/target. These normalized samples were then analyzed with t-SNE (Maaten, 2014). The following parameters were used in the t-SNE analyses: perplexity 10, theta 0, iterations 3000. RNA sequencing samples in the SMARCB1 re-expression studies were subsequently aligned and analyzed with the Tuxedo suite (e.g. aligned with TopHat 2.0.11, abundance estimation with CuffLinks, differential analysis with CuffDiff and CummeRbund) (Trapnell et al., 2010). For the comparison to previously published work (Wang et al., 2016), the published RNA sequencing samples along with our samples were re-aligned with TopHat 2.0.11 and analyzed with the Tuxedo suite. For samples treated with DMSO or MLN2238, samples were aligned as above and analyzed with DESeq2 (Love et al., 2014).

### Sanger sequencing confirmation of WGS findings

gDNA was extracted using QIAamp DNA mini kit (Qiagen). We performed a mixing study of our RMC cell lines with gDNA isolated from the G401 MRT cell line and then performed PCR amplification. We determined that the lower limits of detection of these fusions with our methods were ~1% of tumor cell line gDNA with a minimum 50ng of gDNA (**Supp Fig. 1E**). We subsequently performed the same PCR reactions with 100ng of gDNA from the tumor tissue samples. Samples were gel purified and submitted for Sanger sequencing (Eton Bio). We found that the sequences from the tumor tissue samples matched those of the cell lines (**Supp Fig. 1F and G**), confirming that the genomic alterations that we found in the cell lines reflect those found in the original tumor. Primers utilized were CLF_PEDS0005 *chr 1* forward (ATAAGACATAACTTGGCCGG), CLF_PEDS0005 *SMARCB1* reverse (TTTTCCAAAAGGTTTACAAGGC), CLF_PEDS9001 *chr12* forward (AAAAGCATATGTATCCCTTGCT), CLF_PEDS9001 *SMARCB1* reverse (CCTCCAGAGCCAGCAGA).

### Quantitative RT-PCR

RNA was extracted as above and normalized using the Nanodrop to 1ug. One ug of RNA was then added to the High Capacity cDNA Reverse Transcription Kit (Thermo Fisher Scientific) and PCR reactions were performed as per manufacturer’s recommendations. cDNA was then diluted and added to primers (**Supp Data 11**) and Power SYBR Green PCR Master Mix (Thermo Fisher Scientific). Samples were run on a BioRad CFX384 qPCR System in a minimum of technical quadruplicates. Results shown are representative of at least 2 biological replicates.

### Gene-Expression Array analysis

We performed Affymetrix Human Genome U133 Plus 2.0 on our RMC cell lines. We then combined the following GEO datasets using a GenePattern module with robust multi-array (RMA) normalization GSE64019, GSE70421, GSE70678, GSE36133, GSE94321 (Barretina et al., 2012; Calderaro et al., 2016; Johann et al., 2016; Richer et al., 2017; Wang et al., 2016). We utilized COMBAT and then tSNE to account for batch effects and to identify clusters of similarity (Chen et al., 2011; Johnson et al., 2007; Maaten, 2014).

### Glycerol Gradients followed by SDS-PAGE

Nuclear extracts and gradients were performed as previously published(Boulay et al., 2017). Briefly, 500ug of nuclear extract from approximately 30 million cells was resuspended in 0% glycerol HEMG buffer containing 1mM DTT, cOmplete protease inhibitors and PhosStop (Roche). This was placed on a 10-30% glycerol gradient and ultracentrifuged at 40k RPM for 16 hours at 4C. Following centrifugation, fractions of 550uL were collected. Samples were then prepared with 1x LDS Sample Buffer (Thermo). Samples were run on a 4-12% Bis-Tris gel and then transferred by immunoblotting in tris-glycine-SDS buffer with methanol. Immunoblots were subsequently blocked with Licor Blocking Buffer (Lincoln, NE) and then incubated with antibodies as noted in the methods section. Immunoblots shown are representative of at least 2 biological replicates.

### Cell lines

Primary cell lines were authenticated by Fluidigm or WES/WGS sequencing and tested for mouse antibody production and mycoplasma. Established cell lines were authenticated by Fluidigm SNP testing. Cell lines were refreshed after approximately 20 passages from the frozen stock.

### Small molecules

Bortezomib or MLN2238 was purchased from SelleckChem (Houston, TX) for the *in vitro* studies. Compounds were resuspended in DMSO and frozen down in 20uL aliquots to limit freeze-thaw cycles. Compounds were added as noted in the figure legends. *In vitro* studies used 15nM for bortezomib and 100nM for MLN2238. For the pulse experiments, we used 2.5uM of MLN2238.

### SMARCB1 induction studies

pDONR223 SMARCB1 was Sanger sequenced (Eton Bio) and aligned to the variant 2 of SMARCB1. SMARCB1 was subsequently cloned into the inducible vector pLXI401 or pLXI403 (Genomics Perturbation Platform at the Broad Institute, Cambridge, MA) by Gateway Cloning. LacZ was used as a control. Lentivirus was produced using tet-free serum (Clontech, Mountain View, CA). Cell lines were infected with lentivirus to generate stable cell lines. Cell lines were then confirmed to re-express LacZ or SMARCB1 by titrating levels of doxycycline (Clontech). Parental cell lines were treated with increasing doses of doxycycline to determine the toxicity to cells and measured by Cell-TiterGlo after 96 hours or by trypan blue exclusion cell counting after 2 weeks (**Supp Fig. 2A-B**). Cells were then grown with or without doxycycline in a 6 well plate. Cells were counted by Trypan blue exclusion on a ViCELL XR (Beckman Coulter, Brea, CA) every 4-5 days. Results shown are the average of at least 3 biological replicates.

### Scenesence assays

Cells were plated in a 6 well dish and treated with or without doxycycline for up to 7 days. Senescence was assessed with the Senescence β-Galactosidase Staining Kit without modifications (Cell Signaling Technologies, Danvers, MA).

### Druggable Cancer Targets (DCT) v1.0 shRNA/sgRNA libraries, pooled screens and small-molecule profiling

These were performed as previously published (Hong et al., 2016). Briefly, we utilized the DCT v1.0 shRNA (CP1050) and sgRNA (CP0026) libraries from the Broad Institute Genetic Perturbation Platform (GPP) (http://www.broadinstitute.org/rnai/public/). Viruses from both pools were generated as outlined at the GPP portal. As CLF_PEDS0005_T2A and CLF_PEDS0005_T2B were expanded, we performed titrations with the libraries as outlined at the GPP portal. For the sgRNA pool, both cell lines were first transduced with Cas9 expression vector pXPR_BRD111. We screened the DCT v1.0 shRNA library in biological replicates and the Cas9 expressing cell lines with the sgRNA pools at an early passage (<20) and at a multiplicity of infection (MOI) <1, at a mean representation rate above 500 cells per sgRNA or shRNA. gDNA was extracted and was submitted for sequencing of the barcodes. We achieved sequencing depths of at least 500 reads per shRNA or sgRNA.

### CRISPR-Cas9 validation studies

sgRNAs targeting the genes noted in the manuscript (e.g. PSMB5, UBE2C and controls; **Supp Data 12**) were generated and introduced into the pXPR_BRD003 backbone. These were then sequence confirmed by Sanger sequencing (Eton Biosciences). Lentivirus was produced and used for infection to generate stable cell lines expressing Cas9. Cells were counted or harvested for protein as noted in the text.

### Immunoblots

After indicated treatments, cell lysates were harvested using RIPA buffer (Cell Signaling Technologies) with protease inhibitors (cOmplete, Roche) and phosphatase inhibitors (PhosSTOP, Roche). Antibodies used were as follows: ARID1A (Santa Cruz; sc-373784), α-tubulin (Santa Cruz; sc-5286), β-Actin (C-4) (Santa Cruz; sc-47778), β-Actin (Cell Signaling; 8457), BAF155 (Santa Cruz; sc-10756), BAF170 (Santa Cruz; sc-166237), SMARCA4 (Santa Cruz; 17798), BRM (Bethyl; A301-015A), CDT1 (Cell Signaling; 8064S), Cleaved Caspase-3 (Cell Signaling; 9664), c-MYC (Santa Cruz; sc-764) or c-MYC (Cell Signaling; 9402), cyclin B1 (Cell Signaling; 4135 and 4138), cyclin D1 (Santa Cruz; sc-718), Flag M2 (Sigma; F1804), GAPDH (Cell Signaling; 2118S and 97166S), GRP78 (Rockland Antibodies, Limerick, PA; 200-301-F36), IRE1-alpha (Cell Signaling; 3294), lamin A/C (Cell Signaling; 2032), PSMB5 (Abcam, Cambridge, MA; ab3330), p21 Waf1/Cip1 (Cell Signaling; 2947), p53 (DO-1) (Santa Cruz; sc-126), SMARCB1/SNF5 (Bethyl A301-087A), SPARC (Cell Signaling; 5420S), Ubiquitin (Cell Signaling; 3936), UBE2C (Proteintech, Rosemont, IL; 66087-1). Results shown are representative of at least 2 biological replicates.

### Cell cycle and Annexin V/PI

Cells were treated using the conditions noted in the text. Two million cells were spun down and resuspended in PBS. Cells were then subjected to FITC Annexin V and PI staining as described (BD Pharmigen; 556547). Another set of cells were subjected to PI/RNAse staining (BD Pharmingen; 550825 or Invitrogen F10797). Samples were analyzed within 1 hour with the SA3800 Spectral Analyzer (Sony Biotechnology). Biological replicates were performed. Data was analyzed with FlowJo v10 (FlowJo, Ashland, OR).

### Proteasome function assay

We measured the cell’s ability to cleave Suc-LLVY-aminoluciferin utilizing Proteasome-Glo (Promega) following a one-hour treatment with the noted proteasome inhibitor and measured luminescence. Results shown are from at least 2 biological replicates.

### *In vivo* tumor injections

This research project has been reviewed and approved by the Dana-Farber Cancer Institute’s Animal Care and Use Committee (IACUC), in compliance with the Animal Welfare Act and the Office of Laboratory Welfare (OLAW) of the National Institutes of Health (NIH). Five million cells of G401 in 100μL of a 50% PBS / 50% Matrigel (BD Biosciences) mixture were injected subcutaneously into flanks unilaterally in Taconic NCr-Nude (CrTac:NCr-Foxn1nu) female mice at 7 weeks of age. When tumors reached approximately 150 mm^3^, mice were randomized into various treatment groups: vehicle control (5% 2-hydroxypropyl-beta-cyclodextrin (HPbCD)) or MLN2238 (7 mg/kg IV twice a week for 4 weeks). MLN2238 (diluted in 5% 2-HPbCD) was purchased from MedChem Express. Randomizations to the treatment arm occurred. Blinding was not performed. Statistical analysis was performed using the two-tailed t-test or Mantel-Cox as noted in the text.

## Supporting information

## Supplementary Materials

**Supp Fig.1:**
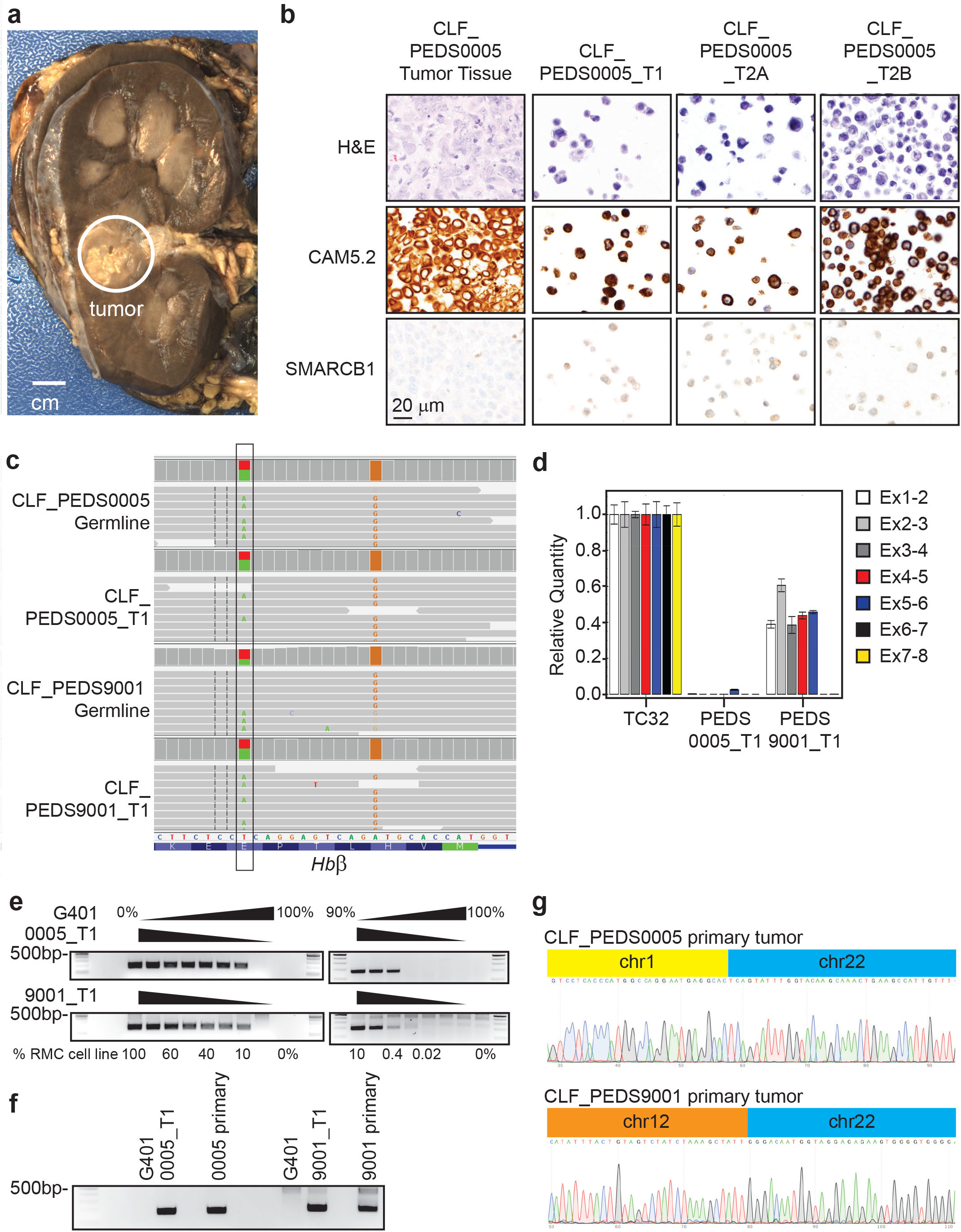
Patient derived RMC models are reflective of known biology and are faithful models of the primary tumors. **(a)** Gross pathology of CLF_PEDS0005 nephrectomy. Circle highlights treated tumor. **(b)** Immunohistochemistry of CLF_PEDS0005 cells highlighting same staining patterns as seen in RMC. **(c)** Integrated Genomic Viewer (IGV) screen shot of the hemoglobin beta loci. At amino acid 7, there is a T/A heterozygous mutation (Glutamic acid/Valine) leading to sickle cell trait. **(d)** Quantitative reverse transcriptase PCR (qRT-PCR) of each exon-exon junction of *SMARCB1* confirms expression loss in RMC samples. Relative expression of these exon-exon junctions in the RMC models was compared to TC32, a Ewing Sarcoma cell line with wild-type *SMARCB1.* **(e)** Mixing studies identify that PCRs are specific to detect genomic fusions between 0.4-2% tumor purity. PCRs spanning the genomic fusion in *SMARCB1* and partner loci were amplified and resolved on 2% agarose gels. Samples were mixed with G401, which has biallelic deletion of SMARCB1, to determine the tumor purity required to identify the fusion sequence. **(f)** PCR amplification of primary tumor samples which were <20% pure identifies *SMARCB1* fusions as identified on PCR-free WGS. **(g)** Sanger sequencing was performed from gel purified products in **(F)** and aligned with PCR-free WGS sequence.

**Supp Fig.2:**
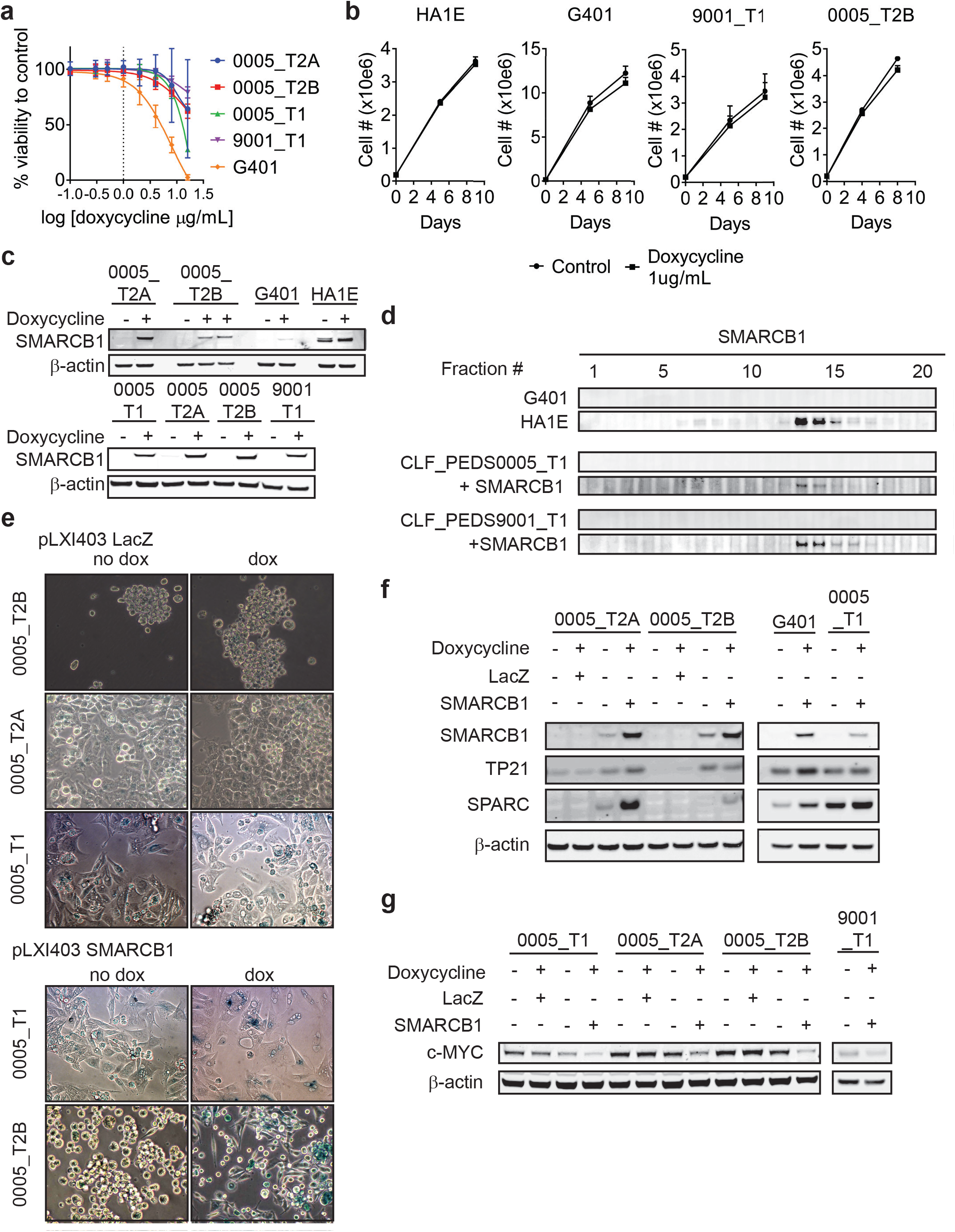
Re-expression of SMARCB1 in patient derived RMC models. **(a)** CellTiter-Glo of parental cell lines treated with increasing doses of doxycycline for 96 hours. Error bars shown are standard deviations from two biological replicates. Dotted line at doxycycline of 1ug/mL (or log 0) indicates dosage used in this study. **(b)** Effect of continuous doxycycline of 1ug/mL over 8 to 9 days. Cells were plated on day 0 and treated with 1ug/mL of doxycycline. Cells were counted at day 4 or 5 and then replated at the same cell number and treated with doxycycline. Error bars shown are standard deviations from two biological replicates. **(c)** SMARCB1 inducible vectors were stably transfected into cell lines. Administration of doxycycline caused similar induction of SMARCB1 across all cell lines. Blots are representative of at least two biological replicates. **(d)** SMARCB1 immunoblot from 10-30% glycerol gradients. SMARCB1 is prominently in fractions 13 and 14. Blots are representative of at least two biological replicates. **(e)** Representative images from three biological replicates of β-galactosidase staining following 7 days of SMARCB1 or LacZ re-expression across CLF_PEDS0005 cell lines. **(f)** Utilizing stably infected inducible cell lines with either LacZ control or SMARCB1, immunoblots performed to confirm transcriptome findings. Following SMARCB1 expression, genes such as TP21 and SPARC were enriched. Blots are representative of at least two biological replicates. **(g)** Utilizing stably infected inducible cell lines with either LacZ control or SMARCB1, c-Myc protein expression is decreased following SMARCB1 expression. Blots are representative of at least two biological replicates.

**Supp Fig.3:**
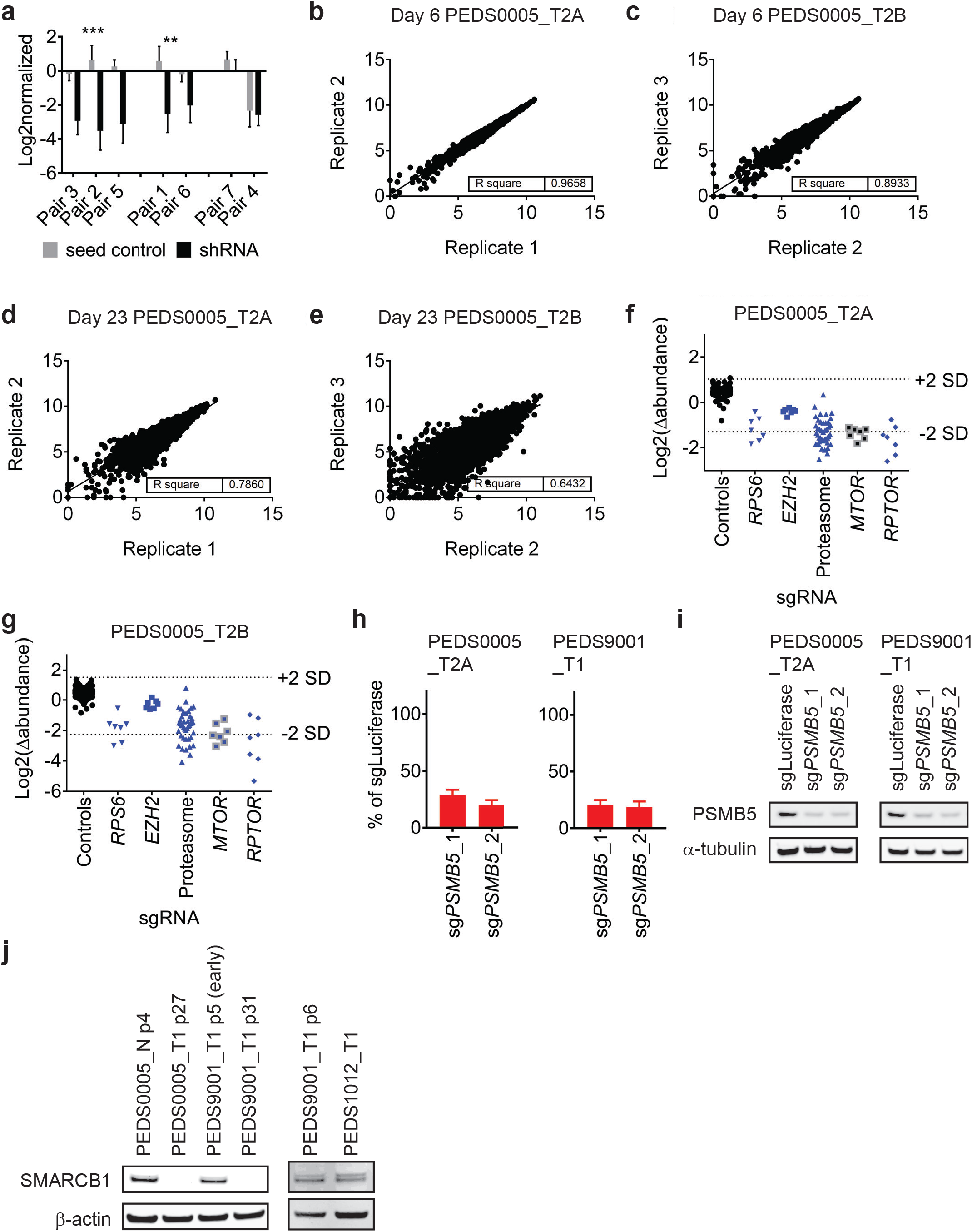
Functional genomic screens identify proteasome inhibition as a vulnerability in RMC and other SMARCB1 deficient cancers. **(a)** Suppression of *RPS6* in CLF_PEDS0005_T2A in RNAi screens. Following log2 normalized counts, shRNAs (black) and paired seed controls (grey) were assessed for off-target effects. Most pairs showed minimal off-target effects while Pair 4 showed significant off-target effects. Error bars represent standard deviation from at least 2 replicates. *** indicates a two-tailed t-test p-value <0.0005, ** <0.005. **(b)** and **(c)** Correlation of replicates from normalized counts of CRISPR-Cas9 screens in CLF_PEDS0005_T2A or CLF_PEDS0005_T2B at the early day 6 timepoint. **(d)** and **(e)** Correlation of replicates from normalized counts of CRISPR-Cas9 screens in CLF_PEDS0005_T2A or CLF_PEDS0005_T2B at the end of the screen (e.g. day 23) timepoint. **(f)** and **(g)** Log2 fold change in abundance of sgRNAs in CRISPR-Cas9 loss of function screens. Controls include sgControls (black) and common essential gene, *RPS6.* In comparison to these, genes involving the proteasome were similarly depleted like *RPS6.* **(h)** Gene deletion by CRISPR-Cas9 of *PSMB5* leads to significant decrease in viable cells in RMC cell lines. Error bars represent standard deviation from at least 2 replicates. **(i)** Gene deletion by CRISPR-Cas9 of *PSMB5* is confirmed by immunoblot. **(j)** Confirmation that the normal cell line, CLF_PEDS0005_N, early passaged tumor cell line, CLF_PEDS9001_T1 and Wilms tumor cell line, CLF_PEDS1012_T 1, express SMARCB1 as compared to the primary RMC cancer cell lines.

**Supp Fig.4:**
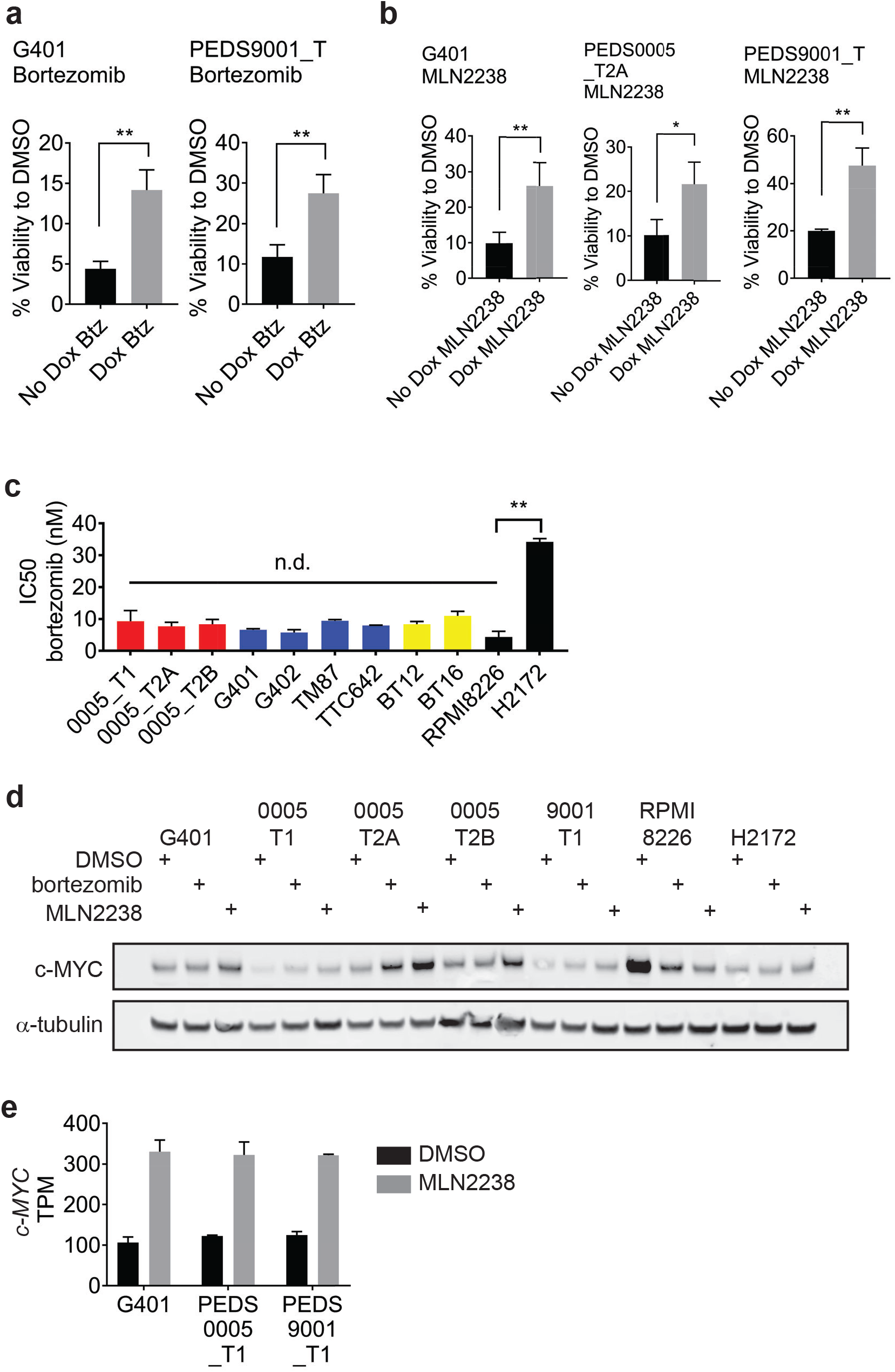
SMARCB1 deficient cell lines are sensitive to proteasome inhibition. **(a)** Using the inducible SMARCB1 cell lines, SMARCB1 deficient lines have increased sensitivity to bortezomib when compared to cells with SMARCB1 re-expressed. Error bars represent standard deviation from at least 3 biological replicates. ** indicates a two-tailed t-test p-value < 0.005 **(b)** Using the inducible SMARCB1 cell lines, SMARCB1 deficient lines have increased sensitivity to MLN2238 when compared to cells with SMARCB1 re-expressed. Error bars represent standard deviation from at least 3 biological replicates. * indicates a two-tailed t-test p-value < 0.05 and ** < 0.005 **(c)** Sensitivity to bortezomib as measured by IC50s. RMC cell lines (red), MRT cell lines (blue) and ATRT cell lines (yellow) are similarly sensitive to RPMI8226, multiple myeloma cell line. This is in comparison to H2172 which is significantly less sensitive to proteasome inhibition. Error bars represent standard deviation from at least 3 biological replicates. n.d. no difference. ** indicates a two-tailed t-test p-value < 0.005. **(d)** c-MYC is not downregulated upon proteasome inhibition at the protein level. Cell lines were treated with DMSO, bortezomib or MLN2238 for 48 hours. c-MYC protein levels in RPMI8226, a multiple myeloma cell line, decrease following proteasome inhibition by immunoblot. However, this does not occur in the SMARCB1 deficient cancer cell lines. Immunoblots are representative of at least 2 biological replicates. **(e)** *c-MYC* is not downregulated upon proteasome inhibition in the transcriptome. Cell lines were treated with DMSO or MLN2238 for 48 hours. Samples from three biological replicates were subjected to RNA-sequencing and c-MYC levels were assessed. Across all the SMARCB1 deficient cell lines, *c-MYC* transcript levels were increased. Error bars represent standard deviation from at least 3 biological replicates.

**Supp Fig.5:**
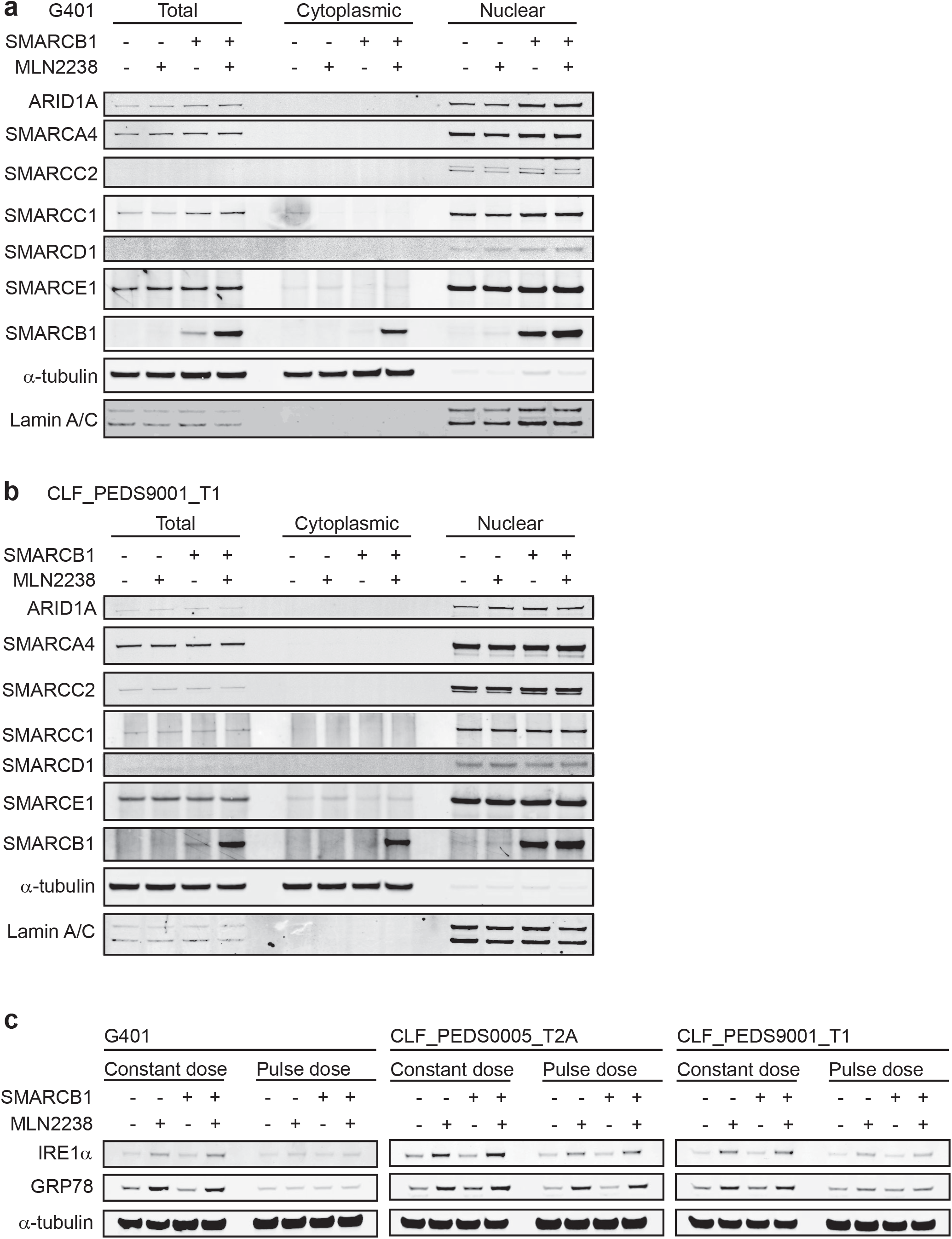
Treatment with proteasome inhibitors does not alter total or nuclear protein levels of SWI/SNF complex members. **(a)** and **(b)** Immunoblots from total, cytoplasmic and nuclear protein fractions show no significant difference in SWI/SNF complex members upon treatment of proteasome inhibitors. Lamin A/C and alpha tubulin are loading controls. Blots representative of at least 2 biological repeats. **(c)** Treatment with proteasome inhibitor, MLN2238, leads to induction of IRE1α and GRP78. However, induction of these ER stress proteins is not rescued upon re-expression of SMARCB1.

**Supp Fig.6:**
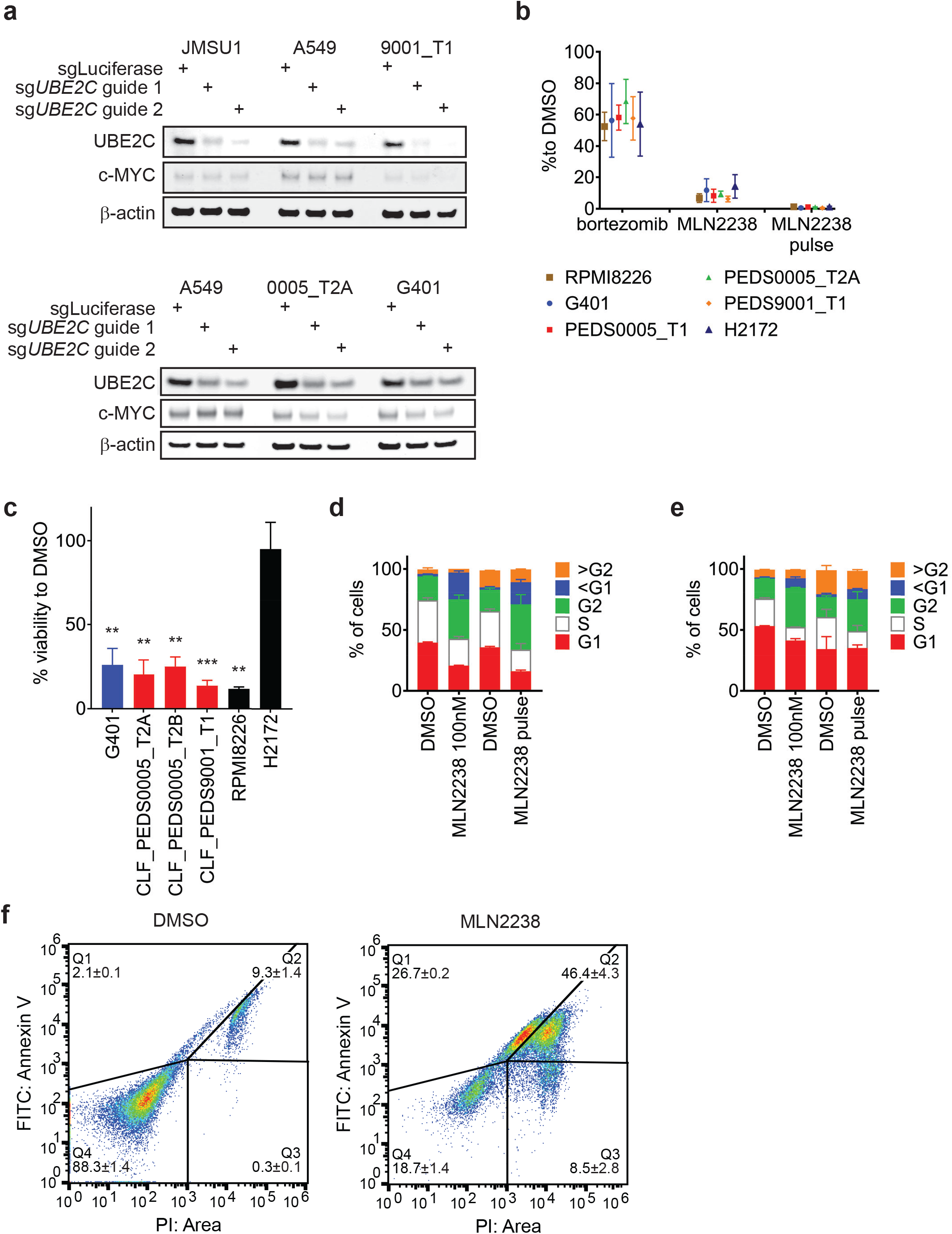
Pulse dosing of proteasome inhibitors leads to suppression of the proteasome, arrests cells in G2/M and leads to programmed cell death. **(a)** Suppression of UBE2C protein by immunoblot when utilizing CRISPR-Cas9 guide RNAs targeting *UBE2C.* cMYC levels modestly decreased upon UBE2C deletion in SMARCB1 deficient cell lines. Blots representative of at least 2 biological repeats. **(b)** Proteasome inhibitors suppress proteasome activity following one hour of treatment by assessing the cell’s ability to cleave Suc-LLVY-aminoluciferin following a one-hour treatment with a proteasome inhibitor as indicated in the figure (**Methods**). Error bars represent standard deviation from at least 3 biological replicates. **(c)** Pulse treatment with MLN2238 leads to significant viability defects in SMARCB1 deficient cell lines similar to multiple myeloma cell line, RPMI8226, and contrasts to lung non-small cell lung cancer cell line, H2172. Error bars are standard deviations from a two biological replicates. **(d)** and **(e)** Cell cycle analysis of G401 **(D)** or CLF_PEDS9001_T1 **(E)** treated with a continuous dose or a pulse dose of MLN2238 shows an increase in cells arrested in G2/M. Error bars represent standard deviation from at least 2 biological replicates. **(f)** Pulse treatment with MLN2238 in CLF_PEDS9001_T1 leads to increased populations that are Annexin V/PI and PI positive. Figures representative of 3 biological replicates.

**Supp Fig.7:**
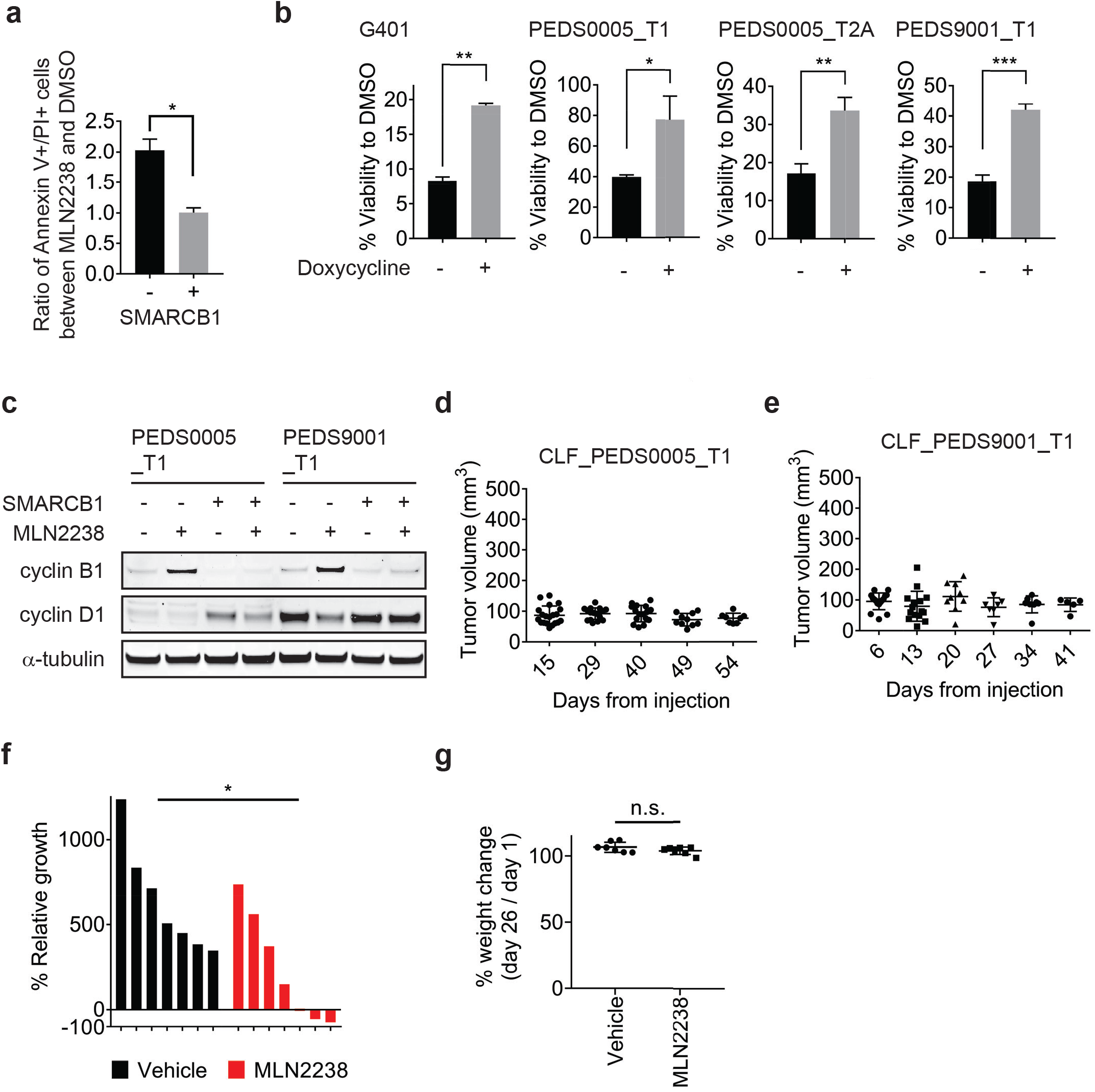
Pulse treatment with MLN2238 can be rescued with SMARCB1 re-expression and in vivo studies identify MLN2238 leading to tumor shrinkage. **(a)** Pulse treatment with MLN2238 in CLF_PEDS9001_T1 in the setting of re-expression of SMARCB1 leads to a decreased fold change in double Annexin V+ / PI+ cells. Error bars represent standard deviation from at least 2 biological replicates. * two-tailed t-test p-value < 0.05. **(b)** Viability defects seen with pulse treatment with MLN2238 can be rescued with re-expression of SMARCB1 in SMARCB1 deficient cell lines. Error bars shown are standard deviations from two biological replicates. * indicates a two-tailed t-test p-value < 0.05, ** <0.005, ***<0.0005. **(c)** Pulse treatment with proteasome inhibitor, MLN2238, leads to upregulation of cyclin B1 and this phenotype is rescued upon SMARCB1 re-expression in both CLF_PEDS0005_T1 and CLF_PEDS9001_T1. Cyclin D1 is included as a control to ensure that the effects of the proteasome are specific to cyclin B1. Blots are representative of two biological replicates. **(d)** and **(e)** Primary tumor RMC cell lines (CLF_PEDS0005_T1 and CLF_PEDS9001_T) do not form tumors *in vivo*. 5 million cells were injected subcutaneously into Taconic immunodeficient mice and were monitored for tumor formation over 41 to 54 days. **(f)** Waterfall plot of relative tumor growth of xenograft G401 tumors at 26 days following treatment initiation with MLN2238. 7 mice accrued in each group were randomized to receive vehicle or MLN2238 once tumors reached an average volume of 148mm^3^. * two-sided t-test p-value <0.05. **(g)** % change in body weight of mice at day 26 as compared to day 1 following treatment with vehicle or with MLN2238. n.s. not significant based on a two-sided t-test p-value.

Supp Data 1. Significant mutations identified by MuTect2

Supp Data 2. SMARCB1 Fluorescence In Situ Hybridization results

Supp Data 3. Structural changes identified by SvABA in CLF_PEDS0005_T

Supp Data 4. Structural changes identified by SvABA in CLF_PEDS9001_T

Supp Data 5. Fusion sequences identified by PCR-Free Whole Genome Sequencing

Supp Data 6. Average differential expression across inducible SMARCB1 RMC and MRT cell lines following SMARCB1 re-expression

Supp Data 7. Overlap between RNAi, CRISPR-Cas9 and small-molecule screens

Supp Data 8. Gene Ontology Gene Set Enrichment Analysis from SMARCB1 re-expression studies

Supp Data 9. Average differential expression across SMARCB1 RMC and MRT cell lines following DMSO or MLN2238 treatment

Supp Data 10. Gene Ontology Gene Set Enrichment Analysis from cells treated with MLN2238

Supp Data 11. SMARCB1 exon-exon junction qRT-PCR primers

Supp Data 12. sgRNAs used in the CRISPR-Cas9 validation studies

## Acknowledgments

We thank our patients, the patient advocacy foundations and the RMC Alliance. We thank John Daley at the Dana-Farber Cancer Institute Flow Cytometry Core Facility, Anita Hawkins at the Brigham and Women’s Hospital CytoGenomics Core, Zach Herbert at the Molecular Biology Core Facilities, Tamara Mason at the Genomic Platform at the Broad Institute, and the Boston University Microarray Resource. We thank the Hahn lab, Boehm lab, Cichowski lab, Kim Stegmaier, Rosalind Segal, David Kwiatkowski, Seth Alper, Toni Choueiri, Carlos Rodriguez-Galindo, the COG Renal Tumors Committee for critical discussions and/or reading of the manuscript. This work was supported in part by the NCI Cancer Target Discovery and Development Network U01 CA176058 (WCH) and U01 CA217848 (SLS), Katie Moore Foundation (JSB), Merkin Family Foundation (JSB), T32 GM007753 (TPH), T32 GM007226 (TPH), AACR-Conquer Cancer Foundation of ASCO Young Investigator Translational Cancer Research Award (ALH), CureSearch for Children’s Cancer (ALH), NCI T32 CA136432-09 (ALH), NICHD K12 HD052896-08 (ALH), Pedals for Pediatrics (ALH), Friends for Dana Farber (ALH), Alex’s Lemonade Stand Young Investigator Grant (ALH), Boston Children’s Hospital OFD BTREC CDA (ALH), NCI P50CA101942 (ALH), Cure AT/RT (SNC/ALH), Team Path to Cure (ALH) and the Wong Family Award (ALH).

## Author contributions

ALH and WCH designed the study. ALH and WCH wrote the manuscript. ALH, YYT, BDK, WJK, PK AS and RD created the cell line models of RMC. ALH, BDK and MBD performed the pooled RNAi and CRISPR-Cas9 screens. ALH and XY sequenced/analyzed the RNAi and CRISPR-Cas9 screens. YYT performed compound assays and YYT and ALH analyzed the results. CC serves as the patient advocacy foundation representative. ALH, GJS, BDK and WJK performed the glycerol gradients. ALH, JW, MG and GK performed the computational analyses. ALH, BDK, MBD, WJK, TH and JL performed the validation studies. ALH, MT, KL and PG performed the mouse studies. ALH and AT prepared the sequencing libraries and managed the sequencing effort. AC reviewed the histopathology and the immunohistochemistry. JSB, BDC, KAJ, KLL, BM, BVH and DS developed the IRB approved protocols. AW and CC managed the tissue biorepository. ALH, OG, PB, SNC, EAM, DR, SLS, PAC, CWMR, CK, RB, KLL, JSB and WCH supervised the studies. All authors discussed the results and implications and edited the manuscript.

## Data and materials availability

All primary data are available from the authors. Noted plasmids in the text are available through Addgene or the Genomics Perturbations Platform at the Broad Institute of Harvard and MIT. CLF_PEDS0005_T1, CLF_PEDS0005_T2B, CLF_PEDS0005_T2A and CLF_PEDS9001_T1 cell lines are available through the Cancer Cell Line Factory at the Broad Institute of Harvard and MIT.

## Accession codes

Sequencing data reported in this paper (whole-genome sequencing and whole-exome sequencing) has been deposited in the database of Genotypes and Phenotypes (dbGaP) under study accession __pending____ and GEO GSE111787.

